# Algae drive convergent bacterial community assembly when nutrients are scarce

**DOI:** 10.1101/2022.06.27.497809

**Authors:** Kaumudi H. Prabhakara, Seppe Kuehn

## Abstract

The assembly of microbial communities is a complex dynamical process that determines community structure and function. Assembly can be influenced by species-species interactions, invasions, the availability of nutrients, and microbial physiology. The interdependence of these factors presents a challenge for understanding community assembly. For example, species-species interactions can be modulated by the availability of nutrients. Here we sought to understand how nutrient supply modulated interactions to affect the assembly process. We exploited algae in association with complex bacterial consortia from soils as models for ubiquitous phototroph-heterotroph communities that play an important role in global primary production. Studying bacterial communities assembled with and without the alga in environments with varying frequency of nutrient supply allowed us to differentiate the impacts of the algae from nutrient availability on the assembly process. A statistical decomposition of community taxonomic structure revealed that it is possible to separate the effects of biotic (presence of algae) and abiotic (nutrient supply rates) factors on community assembly. We found that when the supply of external nutrients is infrequent, the algae strongly impact bacterial community assembly, driving initially diverse bacterial consortia to converge to a common structure. Analysis of sequencing data revealed that this convergence is largely mediated by algal inhibition of specific bacterial taxa. Conversely, when nutrients are supplied with high frequency, bacterial community assembly is not impacted by the presence of the alga. This study shows that complex phototroph-heterotroph communities can be powerful model systems for understanding the assembly process in a context relevant to the global ecosystem functioning.

## Introduction

Building a predictive understanding of the assembly of communities is a central problem in microbial ecology. The assembly process dictates the structure of communities which ultimately impacts their stability and functional properties such as carbon cycling and primary productivity (*1*, *2*). Community assembly depends on an interplay between species-species interactions (*3*), immigration (*4*), emigration, and abiotic factors (*5*) that modulate the growth and physiology of organisms in a collective. Disentangling the relative impacts of biotic and abiotic factors on the structure of a community is challenging because of the interdependence between these factors and the difficulty of controlling them independently in wild settings.

Recent work using microbial consortia has revealed that the local environment has a strong impact on the assembled community. For example, the assembly of marine bacterial communities on polysaccharide particles shows a conserved structure defined by the metabolic traits of the community members (*6*). Similarly, sequencing studies of natural communities in the ocean show that the presumed metabolic traits of the strains present are correlated with local abiotic factors such as light and nutrient levels (*5*). Further, human gut bacterial consortia display a conserved successional process of specific bacterial taxa (*7*). Collectively, these studies point to the idea that the local environment is a strong determinant of the composition of microbial consortia.

What remains unclear is how to differentiate the contributions of abiotic factors such as nutrient levels, temperature and light intensity from the contributions of biotic factors such as species-species interactions. One context where biotic and abiotic factors can both play an important role is the assembly of communities comprised of photosynthetic microbes (algae, cyanobacteria, diatoms) in association with non-photosynthetic (heterotrophic) bacteria. Phototroph-heterotroph communities are pervasive in soils and aquatic environments, where these two metabolic strategies are often found in tight symbioses (*8*–*12*). Phototroph-heterotroph symbioses typically arise from phototrophs fixing CO_2_ and excreting organic compounds (*13*–*18*) that can be subsequently utilized by heterotrophs. However, this is not the only source of organic carbon for heterotrophic bacteria, because they can also utilize nutrients from decaying organic matter. In this sense, the assembly of bacterial communities in the presence of phototorphs is influenced by biotic interactions with the phototrophs and the abiotic availability of nutrients. In fact, studies of wild communities have shown that dissolved organic carbon supplied from terrestrial sources (allochthonous) or excreted from autotrophs (autochthonous) can impact the bacterial community taxonomically (*19*,*20*), functionally (*21*), and dynamically (*22*). These studies point to the idea that environmental variables, such as the availability of nutrients, influence assembly by modulating the relative importance of nutrient exchanges. However, in wild communities causality is challenging to establish and interactions between taxa hard to interrogate.

To overcome these challenges, we used model communities comprised of a dominant phototroph interacting with naturally-derived heterotrophic bacterial communities. We develop the model soil-dwelling alga *Chlamydomonas reinhardtii,* and complex soil-derived bacterial consortia, as a model system for understanding phototroph-heterotroph community assembly (*1*). We cultured the alga with these complex communities and varied the frequency with which exogenous nutrients were supplied. We are able to disentangle contributions of abiotic factors (nutrient supply frequency) and biotic factors (the presence of algae) to community assembly. By comparing bacterial community assembly with and without the alga, we show that when nutrients are supplied frequently the presence of algae does not impact bacterial community assembly. However, when nutrients are supplied infrequently the presence of algae strongly influences community assembly. In a regime where nutrients are supplied infrequently, algae drive initially distinct bacterial consortia to converge to a more similar taxonomic composition. We show that this convergence is driven by algal inhibition of specific bacterial taxa in bacterial communities. Direct measurements of interactions are qualitatively consistent with these results.

## Results

### The need for model phototroph-heterotroph communities

Given the importance of phototrophic microbes in primary production and sustaining bacterial communities, it has been proposed that phototrophs might recruit specific bacterial taxa to their local environment to enable productive symbioses (*8*). This thinking has led to the idea that phototrophic microbes might locally control bacterial community assembly. However, studies attempting to address this question have yielded conflicting results. For example, incubations of lake water with three phototrophs showed that the assembled bacterial communities at the class level were tightly correlated with the identity of the autotroph (*23*). Other studies on algae and diatoms have found similar results (*24*–*26*). Conversely, studies have shown that the identity of the phototroph is only weakly associated with the taxonomy of the associated bacterial consortium (*27*). Further, other works showed that bacterial communities in the phycosphere are weakly taxonomically constrained by the identity of the phototroph to contain certain classes of bacteria such as *Roseobacteria* (*28*, *29*). From these studies it remains unclear when and how photosynthetic microbes impact local bacterial community assembly. Here, we addressed this problem empirically by using naturally-derived bacterial communities and the alga *C. reinhardtii*. Leveraging ‘hybrid’ communities comprised of natural consortia with domesticated phototrophs afforded us control over the composition of the communities while retaining some measure of the complexity of wild communities.

### Algal impact on taxonomic composition depends on the rate of nutrient supply

In conditions where externally supplied nutrients are limiting, we expect the compounds secreted by the algae to have a significant effect on the bacteria that can cohabit with the algae. Therefore, in a system consisting of algae and a diverse pool of bacteria, if nutrients are supplied externally at a high enough rate, we expect the algae to have little effect on selecting the bacteria and the converse.

To test this idea, we used soil samples to initiate communities with a diverse pool of bacteria. To maintain uniformity in algae across the different soils, we used a lab strain of the soil-dwelling alga *C. reinhardtii* (*30*) as the phototroph. We limited the amount of organic carbon in our media (Methods) to enforce interactions between *C. reinhardtii* and the bacterial pool, and added excess amounts of nitrogen (ammonia) and phosphorous to ensure no limitation occurred for these nutrients. To test our hypothesis, we performed serial dilutions on the communities after allowing them to grow for different growth periods - 3 days (*τ*_3_), 6 days (*τ*_6_), 9 days (*τ*_9_), and 12 days (*τ*_12_) (Figure 1). In these conditions, algae reach a saturating density of about 10^7^ cells/mL over a period of about 3 days of growth (inferred from Figure S15). As a result, in the longer growth period conditions, the algae reach saturation after about 3 days, and remain at high density for the rest of the growth cycle. Since algae continue to excrete nutrients even after reaching saturation (*15*), we expect the impact of *C. reinhardtii* to be stronger in communities undergoing dilution every 12 days and weaker in communities undergoing dilution every 3 days. After they are grown for their respective growth periods, the communities were diluted into media with fresh nutrient supply. We performed 10 rounds of serial dilution for each growth period. We used two soil samples (designated “A” and “B”, Methods) to start the experiments. For each growth period, we had five replicates of communities of each soil sample with *C. reinhardtii* (hybrid communities) and two (for soil sample B) or three (for soil sample A) communities without *C. reinhardtii*, as controls (control communities). The communities were open to the atmosphere through sterile foam stoppers to ensure availability of carbon dioxide and oxygen, grown at 30°C with constant stirring, and illuminated from below with cycles of 12 h of light and darkness to mimic day-night cycles (Methods). A schematic of the experimental design is presented in Figure 1.

**Figure 1:**
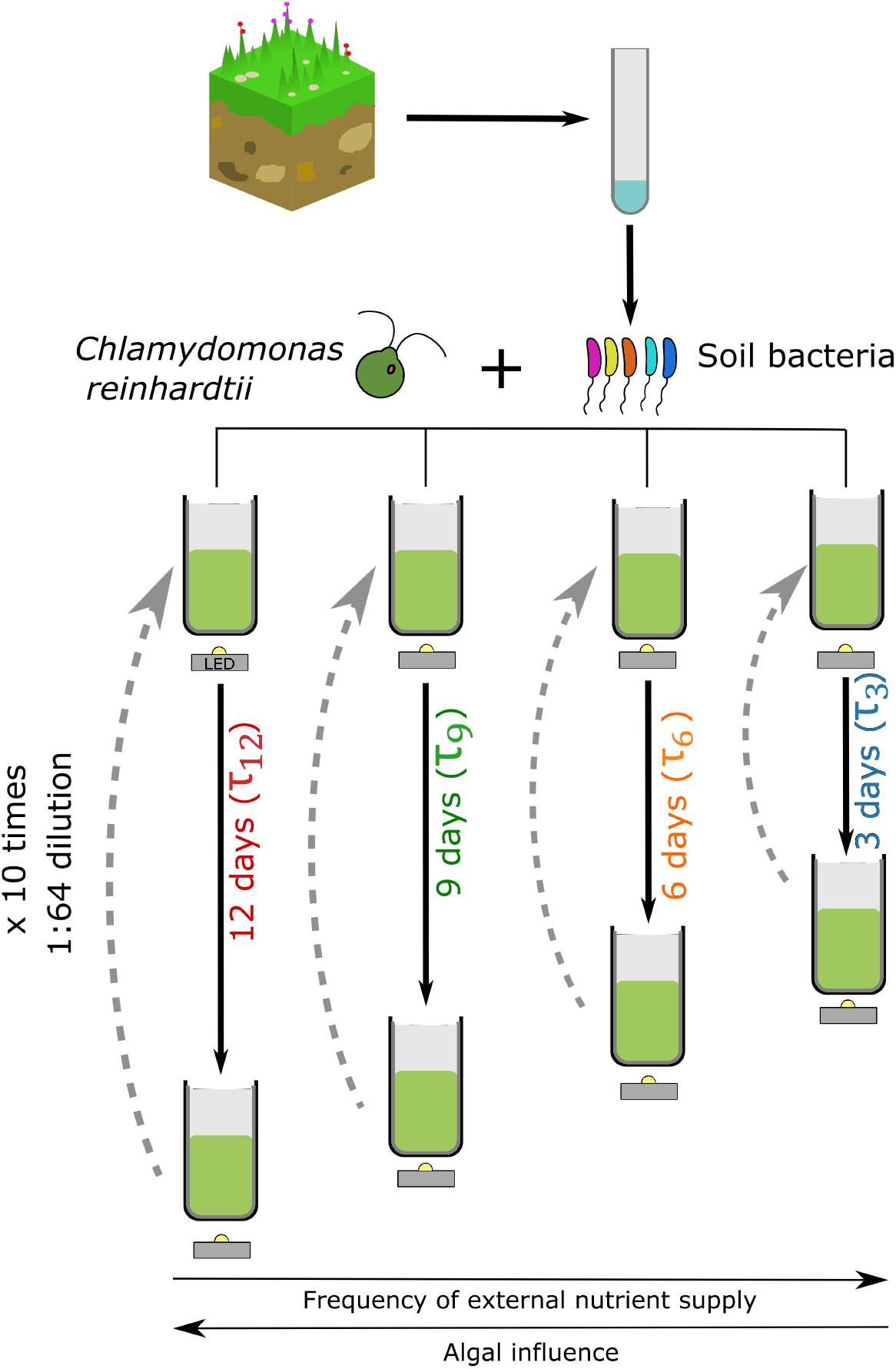
Variable frequency of nutrient supply: Soil samples collected from a restored prairie are treated with drugs to remove eukaryotes and fungi. The soil bacteria are then mixed with a lab strain of C. *reinhardtii,* UTEX2244, mt+. These co-cultures are illuminated with light/dark (12h/12h) cycles and grown for different growth periods - 3 days, 6 days, 9 days and 12 days as indicated. At the end of the growth period, the culture is diluted 64-fold into fresh media. Ten such serial dilutions are performed. Each growth period has 5 replicate cultures. Each growth period also has control cultures, where *C. reinhardtii* is not added (not shown in figure).

In order to asses the effect of the algae on the assembly of the heterotrophic communities, we performed amplicon sequencing of the V4 region of the 16S small subunit rRNA gene on the communities after every other round of serial dilution (Methods, SI Section 1.8). From these data we identified operational taxonomic units (OTUs) in each community using the DADA2 pipeline (*31*) ( SI section 2.1). In Figure 2A and B we show the dominant taxa, defined as those present in at least 20% relative abundance in any time point or community, for the replicate hybrid and control communities respectively.

**Figure 2:**
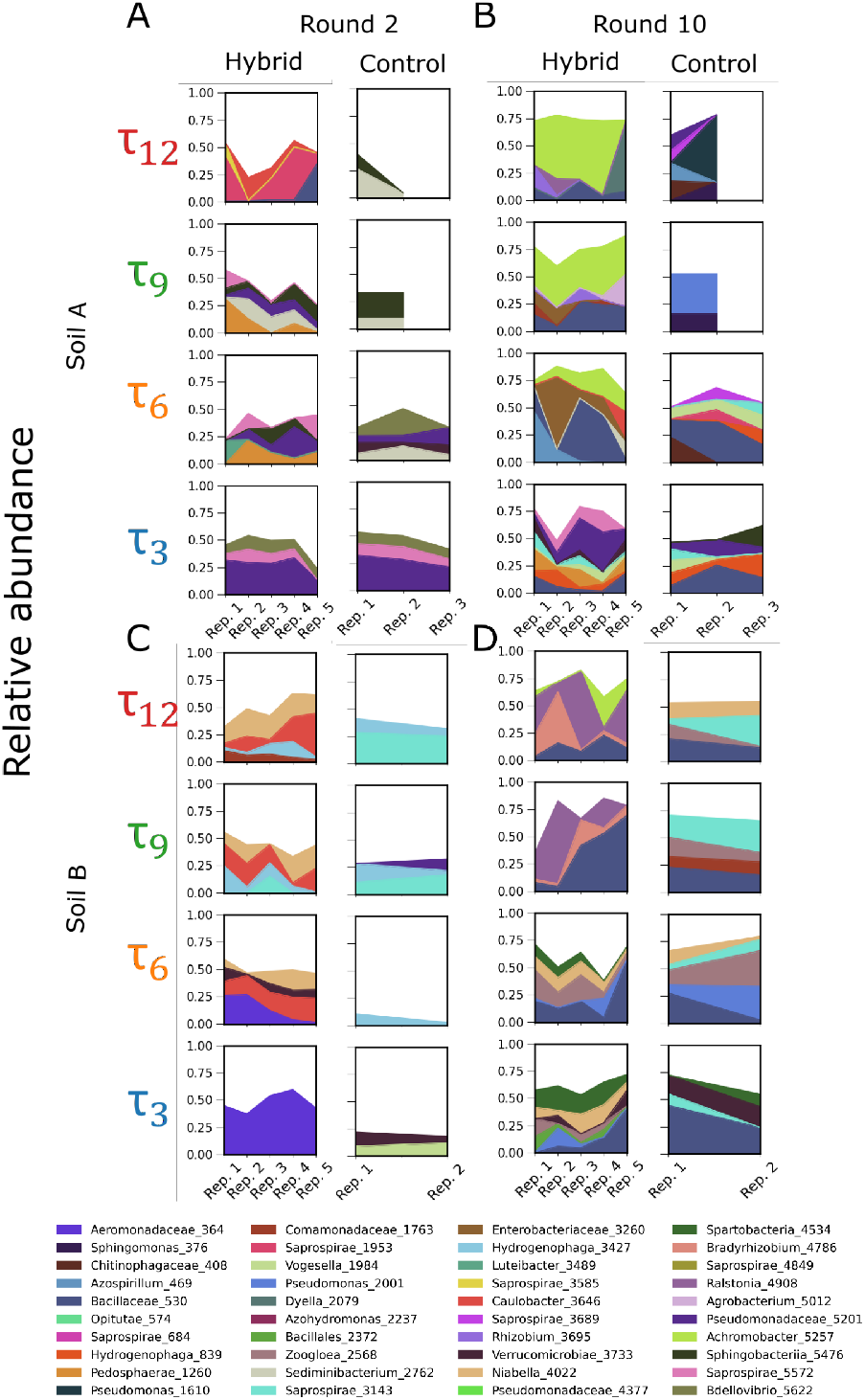
Relative abundance of dominant OTUs at rounds 2 and 10 of serial dilution for hybrid and control communities at all growth periods: Dominant OTUs are defined as those with a relative abundance of at least 20% at any dilution round, soil sample or growth period. Panels A and B show the relative abundance of dominant OTUs in cultures inoculated from soil sample A at rounds 2 and 10 respectively. Rows correspond to growth periods as shown by labels to the left. Panels C and D show dominant OTU relative abundance for cultures inoculated from soil sample B at rounds 2 and 10 respectively. In each panel, the first column shows data for hybrid communities, and the second column shows data for the control communities, as indicated by the labels on the top of each column. Ticks on the abscissa demarcate each replicate community in a given treatment; the hybrid communities have 5 replicates and the control communities have either three (soil A) or two (soil B). See SI for more details. OTU taxonomic classifications are shown in legend at the bottom.

We observe from Figure 2 that in the early dilution rounds, the communities inoculated with different soils are distinct from each other. By dilution round 10, we observe that the hybrid communities inoculated from the same soil have some common dominant OTUs across the 12 day, 9 day and 6 day growth periods within a soil sample (e.g. Ralsontia and Bradyrhizobium in Soil B and Achromobacter in Soil A). However, we do not observe this effect in any *τ*_3_ control communities or the hybrid communities. This suggests that *C. reinhardtii* has a stronger impact on the selection of the OTUs for the longer growth periods, where external nutrient supply is infrequent. The composition of the communities is also distinct from the composition of the soil used to inoculate the communities (Figure S2). The finding that the dominant taxa are similar in hybrid communities with 6, 9 and 12 day growth period holds if we define taxa at the ASV level or at the genus level (Figures S3, S4).

From Figure 2, we also observe that the control communities are taxonomically distinct from the hybrid communities, especially at later dilution rounds. For example, the composition of the *τ*_3_ hybrid and control communities at round 2 for soil A is very similar. This similarity decreases at round 10 (Figure 2). We quantified this similarity using the Jensen Shannon Divergence metric (Figure S5) and found that indeed, the control communities are significantly different from the hybrid communities, and this difference increases with dilution rounds for the longer growth periods (9 and 12 days, Figure S5). This again implies that the presence of *C. reinhardtii* drives the hybrid communities with longer growth periods to be distinct from the control communities. Because the distinction between hybrid and control communities is stronger for the longer growth periods, we infer that the algal impact is stronger in the communities with the longer growth periods, where the frequency of external nutrients supplied is infrequent.

To more globally assess the effect of *C. reinhardtii* on bacterial community assembly, we computed the Shannon diversity index for all communities in our experiment. We found that for hybrid communities with longer growth periods, the diversity decreased with rounds of dilution (Figure 3, left panel). Similarly, *τ*_3_ hybrid communities had higher diversities than *τ*_12_ and *τ*_9_ hybrid communities by dilution round 10. However, the diversity of the *τ*_3_ hybrid and control communities were not significantly different by round 10 (Figure 3, compare *τ*_3_ left and right panels). This indicates that it is the presence of *C. reinhardtii*, whose impact is more strongly felt in the hybrid communities with longer growth periods, that is responsible for the decline in diversity of the hybrid communities with longer growth periods.

**Figure 3:**
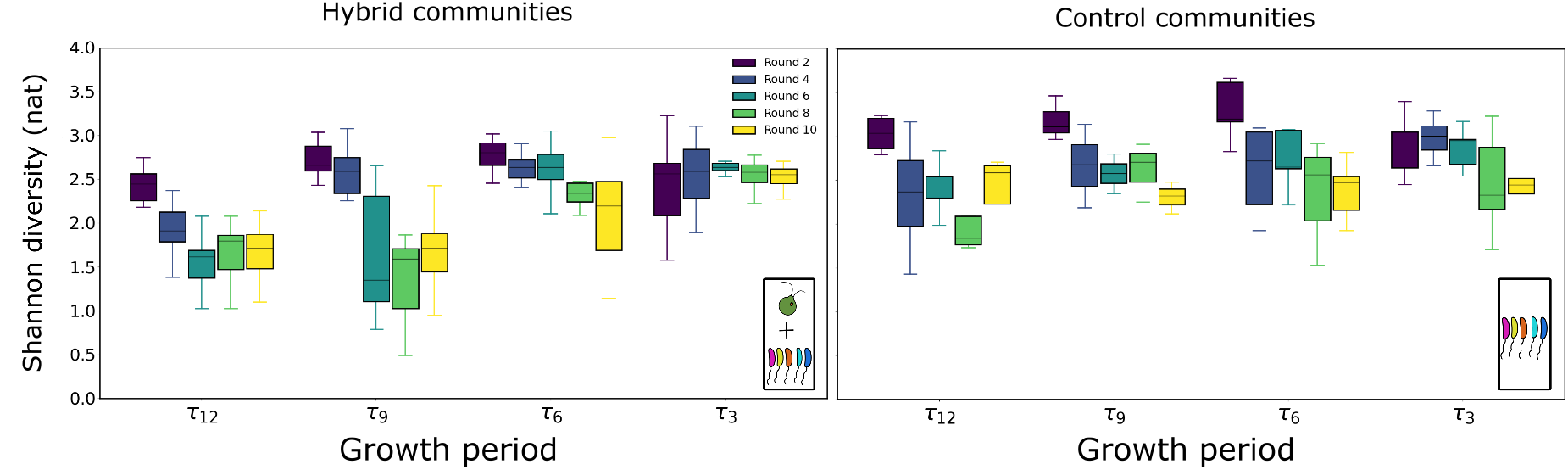
Impact of growth period and presence of algae on community diversity: The distributions of Shannon diversity metrics of the communities across replicates are plotted here as a function of the growth period for the different dilution rounds (black to yellow). The boxes extend from the first to the third quartile, the horizontal line shows the median and the whiskers represent 1.5 times the inter-quartile range (first - third quartile). For the *τ*_9_ and *τ*_12_ hybrid communities, the diversity drops significantly by round 10 compared to their respective control communities at round 10 (p-value 0.05 and 0.03 respectively for the null hypothesis that the diversities of the control communities is the same as the hybrid communities). Similarly, diversity in the *τ*_3_ hybrid communities is higher than *τ*_12_ hybrid communities (p-value of 0.0002 with the null hypothesis that the diversity of the hybrid *τ*_3_ communities is the same as the hybrid *τ*_12_ communities). For shorter growth periods, there is no difference in diversity between control and hybrid communities (p-value of 0.9 for the null hypothesis that *τ*_3_ control communities have the same diversity as the *τ*_3_ hybrid communities.) The Kolmogorov-Smirnov test was used to compute the p-values.

### Dimensionality reduction shows algae drive convergence of bacterial community composition

To see if the impact of algae (biotic factor) and growth period (abiotic factor) on community assembly could be distinguished, we performed principal component analysis (PCA) on the relative abundance data. Specifically, we constructed a data matrix where the rows are communities and the columns OTUs with the entries the number of reads mapping to each OTU in each community. We included all hybrid and control communities, at all growth periods and dilution rounds in this analysis. Data from the initial soil inoculates was also included. The relative abundance data are compositional and therefore must be analyzed with this fact in mind (*32*, *33*). Performing routine statistical analyses on compositional data requires taking log-ratios of the counts data (*34*) after adding pseudocounts to the OTUs sequences receiving zero reads (Methods). Using a permutation approach, we found that only the largest two eigenvalues exceeded the noise in our dataset (Methods). Therefore, we studied the dynamics of our communities along the first two principal components which accounted for 14.6% and 10.4% of the variance of the data respectively.

We present projections of community composition onto the first two principal components in Figure 4. Although a single PCA analysis was performed on all communities at once, for the purpose of visualization we split up the data by the presence/absence of *C. reinhardtii* and growth period. Circles and triangles distinguish the two soil samples and colors distinguish the growth periods. The medium sized translucent markers represent serial dilution round 2, the solid, large markers represent serial dilution round 10, and the smallest markers represent the intermediate rounds. The lines trace the movement of each community across the serial dilution rounds.

**Figure 4:**
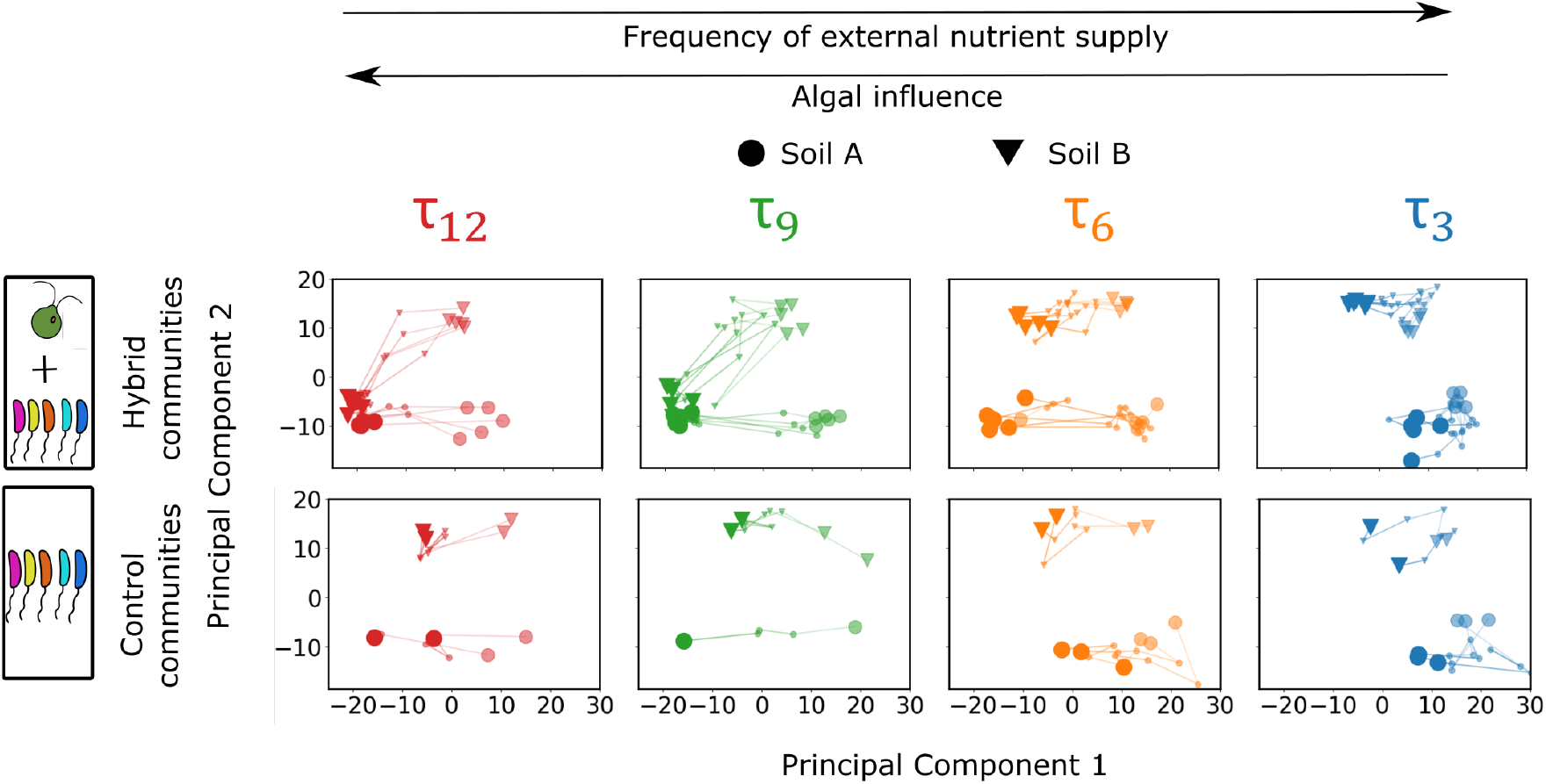
Principal components analysis reveals that the presence of algae mediate community convergence for longer growth periods: Principal components analysis (PCA) was performed on the abundance data for all hybrid and control communities. Rare taxa, defined as those below a threshold relative abundance in all communities and time points, were removed. Removing rare taxa simplifies downstream analysis and does not affect our results (Figure S6). A small number is added to the rest of the data set to remove zeroes and the relative abundances for each community are center-log transformed. PCA was performed on this matrix and two components were deemed significant using a permutation approach (Methods). A single PCA was performed for all communities but for visualization we show growth periods, hybrid, and control communities separately. The top row of panels shows the hybrid communities (those with *C. reinhardtii* as indicated at the left) with growth period decreasing from left to right and the bottom row of panels show the control communities (those without *C. reinhardtii*). In each panel, circles indicate soil A and triangles indicate soil B. The mid-size markers indicate the communities at Round 2 and the largest markers indicate the communities at Round 10. The smallest markers indicate the communities at Rounds 4, 6, and 8. Colors denote growth periods as indicated above each column. Black arrows along the top indicate frequency of external nutrient supply increasing left to right and increasing algal influence on community assembly right to left.

Two main features are clear from Figure 4. First, it is apparent that at round 2, the two soil samples are taxonomically distinct, both in the hybrid and control communities (compare the intermediate sized translucent circles and triangles in all panels of Figure 4). However, with dilution rounds, the *τ*_9_ and *τ*_12_ hybrid communities moved closer to each other, with communities from soil A moving along the first principal component (PC1), and communities from soil B moving along PC1 and principal component 2 (PC2). We refer to the fact that the distance between communities from soil samples A and B rapidly declines with dilution rounds in the *τ*_12_ and *τ*_9_ hybrid conditions as ‘convergence’. Second, the control communities (Figure 4B) do not converge, but they do move along PC1 as do all hybrid communities irrespective of the presence of algae. These features of the community dynamics are also observed when we perform other dimension reduction methods on the data matrix (Figures S9 and S10) or use the data without removing rare taxa (Figure S6C). Next we sought to dissect how the abundance dynamics within the bacterial consortia drove these two outcomes.

### Convergence along principal component 2 arises from algal inhibition of bacteria

Convergence occurred when the *τ*_12_ and *τ*_9_ hybrid communities of soil sample B moved down along PC2 over the rounds of serial dilution to become taxonomically similar to soil sample A by dilution round 10 (Figure 4, *τ*_12_ and *τ*_9_ top row panels). Since this convergence only occurred for the hybrid communities with the long growth periods, the convergence was caused by *C. reinhardtii*. We wanted to understand how the algae affect the heterotrophs over the dilution rounds to cause this convergence. To find the heterotrophs causing motion along PC2, we used the loadings along PC2 to find the contribution of each OTU to the motion along PC2 for the *τ*_12_ and *τ*_9_ hybrid communities of soil sample B. From the distribution of contributions, we selected those OTUs that had the highest contribution (Figure 5 A, SI Section 2.5) to the displacements. These are the OTUs that were the most significant contributors to motion along PC2. Since we have established that *C. reinhardtii* causes the displacement along PC2, we call these bacterial taxa, through which the effect of the algae is manifested, the “biotic taxa”. We found that these sets of taxa for hybrid communities with growth periods *τ*_9_(*S*_*τ*_9__) and *τ*_12_(*S*_*τ*_12__) have more than 90% taxa in common; i.e. C. *reinhardtii* affects the same taxa for both growth periods to displace the communities along PC2 (Figure 5B). In other words, changes in the relative abundances of the same OTUs caused motion along PC2 for both the *τ*_9_ and *τ*_12_ hybrid communities in soil B.

**Figure 5:**
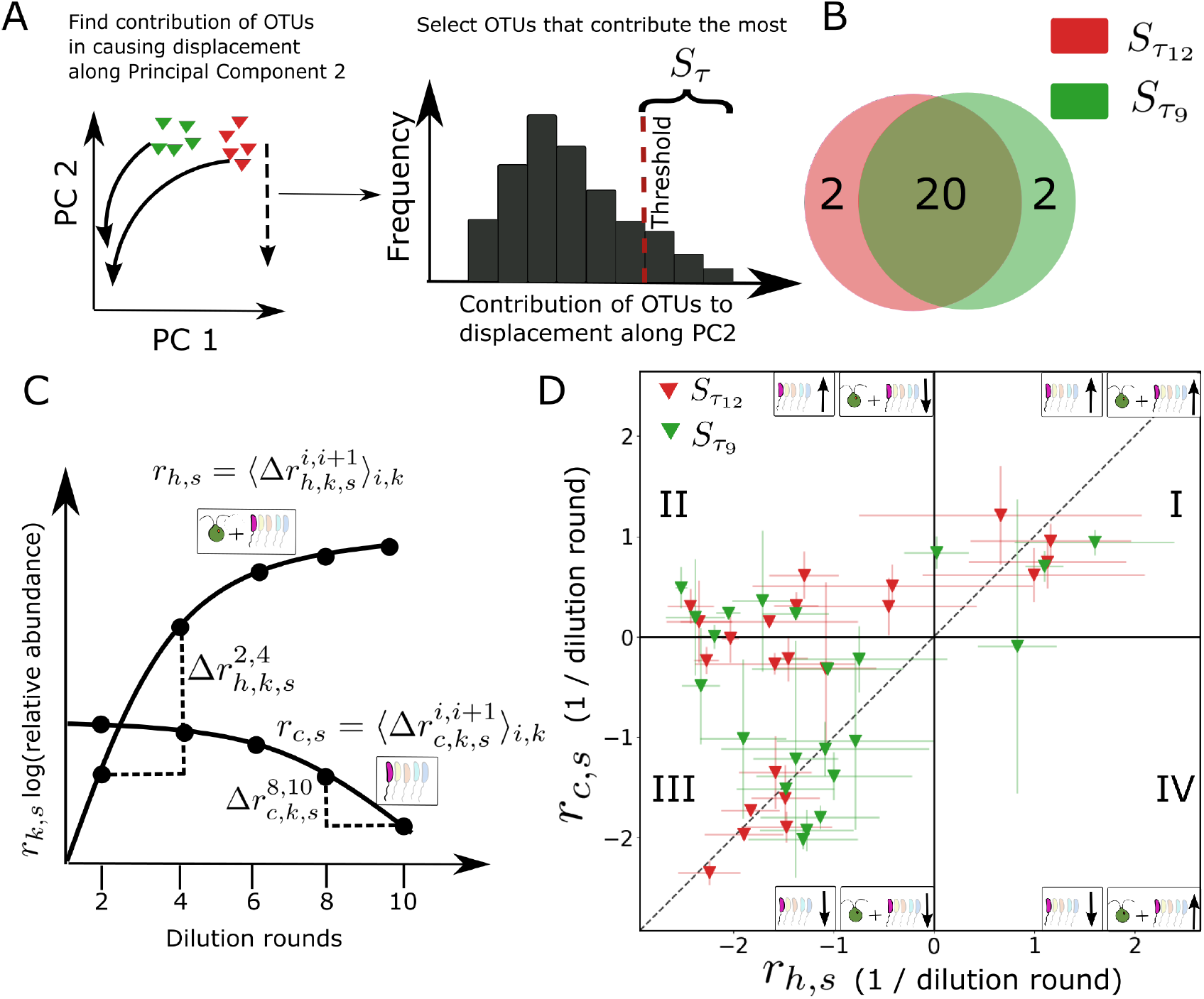
Hybrid community convergence arises from inhibition of bacterial taxa in the presence of algae: A. The contribution each OTU to the displacement along PC2 is found for *τ*_12_ and *τ*_9_ hybrid communities from soil B (triangles). A histogram of the contribution of each taxa to displacement along PC2 is computed and the taxa with contributions above a threshold are selected (*S_τ_*). B. A Venn diagram comparing the identity of taxa selected by the procedure in panel A for both 9 and 12 day growth periods. C. To compare the dynamics over the rounds of serial dilution, enrichment of these biotic taxa is computed in both hybrid and control communities. We denote the logarithm of the relative abundance of OTU *s*, in replicate *k*, at time point *i* for a hybrid community as 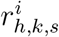. The change in relative abundance between two adjacent time points is then 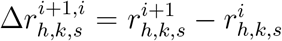. For each OTU we then average these changes across time points and replicates: 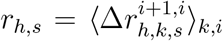. For each OTU we perform the same analysis in control communities to estimate the enrichment in the absence of algae (*r_c,s_*). D. The enrichment rates in the hybrid community (*r_h,s_*) are plotted against the rates in control communities (*r_c,s_*) for each OTU selected in panel A. The error bars indicate standard deviation over replicates. Colors indicate the growth period used to compute the enrichment. See main text for interpretation of each quadrant labeled with roman numerals.

This simple analysis tells us which taxa are driving the convergence along PC2 but does not reveal the dynamics of these taxa over the rounds of serial dilution. Therefore, we sought to understand how these taxa (*S*_*τ*_12__ and *S*_*τ*_9__), that drive convergence in hybrid communities were changing in time over the course of the enrichment experiment. To obtain their dynamics over the dilution rounds, we found the differences in the logarithm of the relative abundances (*r*) of the biotic taxa over the successive dilution rounds in both hybrid and control communities. For a given OTU *s* ∈ *S*_*τ*_12__ or *s* ∈ *S*_*τ*_9__, we found 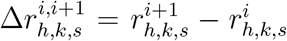, where *h* indicates a hybrid community, *i* represents the round of dilution and *k* the replicate. We average over all time points and replicates (Figure 5C) to find the average net enrichment in the presence of the algae, 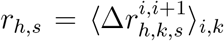. Note that the sign of *r*_h,s_ tells us, on average, whether the relative abundance of OTU *s* is increasing or decreasing over dilution rounds. We also calculate this enrichment for the same taxa in the absence of algae (control communities) which we denote as *r_c,s_* where the subscript *c* refers to the control communities. As sketched in Figure 5C, a given taxa could have different dynamics in the presence and absence of algae.

By plotting *r_h,s_* verses *r_c,s_* we can gain insight into the impact of algae on each biotic taxa *s*, Figure 5D, where each point corresponds to a single taxon from either *S*_*τ*_9__ (green) or *S*_*τ*_12__ (red). The diagonal dashed line represents the case where the enrichment of the taxa is the same in both the hybrid and control communities. Algae has no effect on the enrichment of the taxa on this line because the enrichment is the same irrespective of the presence of algae. In the first quadrant (I), the enrichment is positive for both control and hybrid communities, implying that the taxa in this quadrant increase in relative abundance over dilution rounds in both the control and hybrid communities. In the second quadrant (II), the enrichment is positive for control communities, and negative for hybrid communities. Taxa in this quadrant increase in abundance in the absence of the algae, but decrease in abundance in the presence of algae, indicating that they are preferentially inhibited in the presence of *C. reinhardtii*. In the third quadrant (III), the enrichment is negative for both the hybrid and control communities. The taxa in this quadrant decrease in abundance both in the presence and absence of algae. However, the taxa present above the dashed line are suppressed more strongly in the presence of the algae, because above the dashed line, the enrichment is more negative in the hybrid communities than in control communities. In the fourth quadrant (IV), the enrichment is positive for the hybrid communities and negative for control communities, implying that the taxa present here increase in abundance in the presence of algae.

We observe that most of the biotic taxa (*S*_*τ*_12__ and *S*_*τ*_9__) are in quadrants II, and III, where the presence of algae suppresses the taxa. The taxa that contribute the most to the motion along PC2 dominantly occupy the second quadrant of the enrichment plot, for both the *τ*_12_ and *τ*_9_ hybrid communities, where the presence of algae strongly suppresses their enrichment. In quadrant III, the taxa located above the diagonal are more strongly suppressed in the hybrid communities compared to the control communities. The almost complete absence of taxa in quadrant IV, where enrichment is positive in the hybrid communities but negative in control communities, indicates that it is rare for algae to recruit taxa to drive taxonomic convergence. The dominant effect of the algae is therefore to suppress these biotic taxa. In other words, the presence of *C. reinhardtii* causes these biotic taxa to decrease in abundance over the dilution rounds. This suppression results in the motion of soil B communities along PC2, driving convergence between soil A and B communities at growth periods of 9 and 12 days. Effectively, hybrid communities of soil sample B became more similar to soil sample A by getting rid of some OTUs. It is important to note, however, that we cannot interpret Figure 5D as characterizing direct interactions between algae and any member of the community. Instead, this analysis characterizes the global impact of the presence of algae on the relative abundances of each OTU of interest, but these changes may be mediated by indirect effects or emerge from networks of interactions in the community. We address this problem below using experiments with isolates.

The finding that algae inhibit specific OTUs in soil B to drive convergence with soil sample A makes sense in light of the results in Figures 2, 3, 4 and Figure S1. First, a close examination of the dominant OTUs in *τ*_12_ and *τ*_9_ hybrid communities at round 10 in soil sample A and B reveals that the dominant taxa are not the same between these two communities (Figure 2B, hybrid column). As a result, convergence in Figure 4 *τ*_12_ and *τ*_9_ hybrid communities arises not from algae recruiting similar dominant taxa, but from the inhibition of taxa that are unique to soil B. In addition, the finding that algae preferentially inhibit bacteria in hybrid communities (Figure 5D) is consistent with the loss of diversity in the hybrid communities (Figure 3) at longer growth periods. Further, measurements of optical density (OD) and chlorophyll fluorescence (Figure S1) reveals that the *τ*_3_ hybrid communities have a higher OD per chlorophyll content compared to the *τ*_12_ hybrid communities, indicating that there are more heterotrophs per algae in the *τ*_3_ hybrid communities compared to the *τ*_12_ hybrid communities, consistent with the observed loss of diversity in Figure 3.

In contrast to the long growth period communities, hybrid communities with a 3 day growth period (*τ*_3_) do not converge (Figure 4). We concluded that the longer growth periods drive this convergence, but it is reasonable to ask whether the 3 day hybrid communities simply did not have enough time to converge due to shorter total incubation time: 10 rounds × 3 days - 30 days, versus 120 days in the 12 day growth period experiment. To interrogate this possibility further we first examined the dynamics of the *τ*_3_ hybrid communities from soils A and B. Unlike communities at longer growth periods, we found that the distance between *τ*_3_ hybrid communities from the two soil samples along PC2 increased over dilution rounds, whereas for the 9 day and 12 day growth periods this distance decreased over dilution rounds (Figure S11). Second, we analyzed the dynamics of *S*_*τ*_12__ and *S*_*τ*_9__ taxa in the *τ*_3_ communities using the relative enrichment approach developed above. In contrast to our results in Figure 5D, where the taxa are predominantly on the left half of the plot, where they are suppressed in the presence of *C. reinhardtii,* in the *τ*_3_ and *τ*_6_ communities we found that the biotic taxa are predominantly along the diagonal of the enrichment plot (Figure S12) indicating that *C. reinhardtii* does not impact the bacterial enrichment rates significantly in these conditions. This suggests that the soil A and soil B *τ*_3_ hybrid communities would not converge even given additional rounds of serial dilution. We propose that the reason *τ*_3_ hybrid communities do not converge is because over just 3 days *C. reinhardtii* has a significantly less time to impact bacterial growth. Specifically, we estimated the growth rate of the algae in these conditions to be approximately 8 hours. At this growth rate, it takes the algae approximately 3 days to reach saturation after each round of 64-fold dilution. In contrast, during a 12 day growth period, algae reach saturation after approximately 3 days and remain at high levels for the remaining 9 days. We expect that during this 9 day period algae continue to excrete nutrients and other compounds (*15*) potentially impacting bacterial community assembly. Collectively, these results support the idea that the 3 day hybrid communities are unlikely to converge even with additional rounds of serial dilution.

### Dynamics along PC1 are driven by abiotic factors

From Figure 4 we observe that all communities, irrespective of the presence of the algae, move along PC1. Further, the actual displacement along PC1 depends on the growth period, i.e. the frequency of supply of external nutrients. The *τ*_3_ communities, both hybrid and control, are displaced much less along PC1 compared to the *τ*_12_ communities, both hybrid and control (Figure S13). Because the motion along PC1 is independent of the presence of algae, but depends on the growth period, we hypothesize that the motion along PC1 is caused by abiotic factors, in particular, the frequency of nutrient supply.

To confirm that the presence of algae does not impact the dynamics along PC1, we performed the following test. Using the loadings along PC1, we found the contribution of each OTU to the displacement along PC1 for each community. From these, we selected the major contributors using the cumulative distribution of the contributions (as in Figure 5A, SI Section 2.6). Using this approach, we identified the taxa driving dynamics along PC1 for each community. We performed Fisher’s exact test with the null hypothesis that the taxa contributing to motion along PC1 for a given soil sample and growth period are equally likely to be selected in the control and hybrid communities. We found that we were not able to refute the null hypothesis with a p-value corrected for multiple tests using the Bonferroni correction (See SI Section 2.6 and Dataset 5). We conclude that the same taxa cause motion along PC1 in both the hybrid and control communities, indicating that, statistically, there is no evidence that algae drive motion along PC1. This means that that motion along PC1 is caused by an abiotic factor, the frequency of external nutrient supply, or growth period, common to the hybrid and control communities.

### Algal impact on bacterial isolates varies

Our analysis of relative abundance dynamics in the enrichment experiment suggested that the dominant effect of algae is to suppress bacterial taxa responsible for motion along PC2 in the long growth period conditions (Figure 5D). As discussed above, this measurement does not directly characterize interactions between algae and the associated bacterial taxa. To directly elucidate the types of interactions driving dynamics in our communities we isolated 21 unique bacterial strains from the communities (See SI Section 1.9 and Dataset 8). These strains represented the taxonomic diversity of the assembled communities well, spanning all four quadrants of the enrichment plot (Figure S14). Further, two of these isolates (Bacillacea 530 and Rasltonia 4908) were dominant taxa as captured by our amplicon sequencing measurements (Figure 2).

Since we observed primarily inhibition of the biotic taxa in our enrichment analysis of the sequencing data (Figure 5D), we first wanted to directly assay the effect of algae on bacterial growth. To accomplish this, we grew each isolate on its own and in the presence of algae in conditions identical to the enrichment experiment (Methods). We assayed final bacterial carrying capacity after two weeks of co-culture, quantifying bacterial abundances via measurements of colony forming units (CFUs). To assess the impact of algae on bacteria we compared the carrying capacity of bacteria with and without the algae. We observe that for 5 of the 9 isolated strains there was no significant effect of the algae (Figure 6A). In two cases, the presence of the algae significantly inhibited the growth of the bacterial isolates. Algae significantly enhanced the growth of two other isolates.

**Figure 6:**
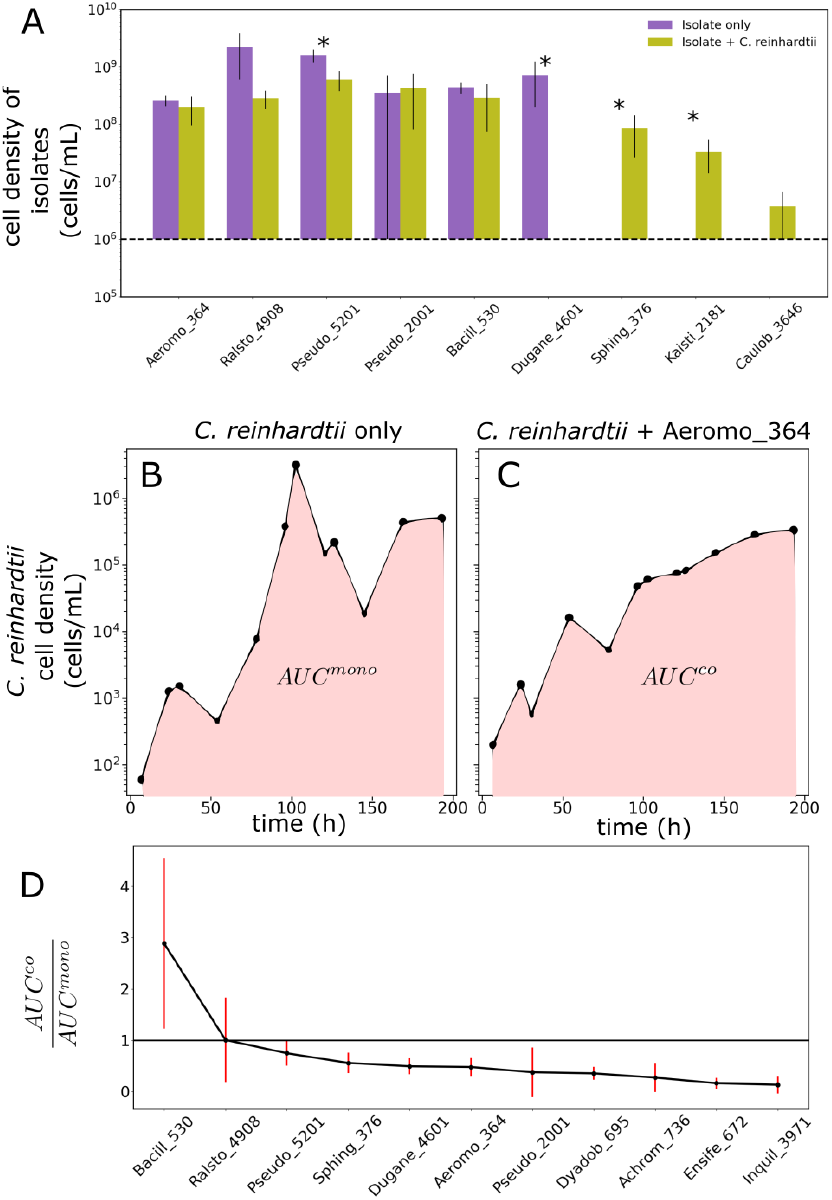
Bacteria inhibit algal growth, but can be either positively, negatively or neutrally affected by algae: Panel A shows the impact of algae on bacterial growth. Panels B-D show the impact of bacteria on algal growth. A Characterizes the impact of algae on the growth of bacteria. Each bacterial isolate was grown in co-culture with *C. reinhardtii* in 2-5 replicates. After 14 days bacterial density was assayed by plating (Methods). Bars indicate measured bacterial density with (yellow) and without (purple) algae. The first six isolates were either inhibited by algae or algae had no impact on growth. The last three bacterial isolates, which grow to a density below the detection limit of 1 × 10^6^ cells/mL (represented by the horizontal dashed line) on glucose, grow better in the presence of algae than in the absence. The stars indicate the cases where growth with and without the algae are significantly different as tested by Kolmogorov Smirnoff test. B shows an example trace of cell density of *C. reinhardtii* grown alone. Densities are measured by flow cytometry. The shaded area represents the area under the curve (*AUC*) for the monoculture, *AUC^mono^*. C shows an example trace of cell density of *C. reinhardtii* grown in co-culture with a bacterial isolate (Aeromo 364). The shaded region represents the AUC for co-culture, *AUC^co^*. D shows the are ratio 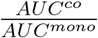 for different bacterial isolates from our communities. This ratio is larger than 1 for bacteria that enhance algal growth relative to monoculture and <1 for bacteria that inhibit algal growth. Each co-culture and the monoculture were run in triplicate, errors in 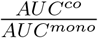 were computed via error propagation from the standard deviation across the three replicates. With the exception of Bacill 530, all other isolates have either a inhibitory or neutral effect on the growth of the algae.

We wanted to check whether our measurements of algal impacts on bacterial growth were consistent with our analysis of the sequencing data. For example, we might naively expect that taxa present in quadrant II of Figure 5D to be inhibited by the algae. Therefore, we performed an identical analysis to Figure 5D for OTUs that correspond to our library of isolates (Figure S14).

Of the 9 bacterial isolates used here, Aeromo 364, Ralsto 4908, Pseudo 5201, Kaisti 2181, and Caulob 3646 belong to the set of biotic taxa (*S*_*τ*_9__ and *S*_*τ*_12__). Aeromo 364 and Ralsto 4908 are close to the diagonal on the enrichment plot (Figure S14), which is consistent with the observation here that the abundances of these two taxa are not significantly impacted by the presence of algae (Figure 6A). Similarly, Pseudo 5201 is located in the third quadrant of the enrichment plot in Figure S14, above the diagonal, implying that the presence of algae inhibits it more strongly than the its absence, consistent with the observation that it is inhibited by the algae. Also, the enhanced growth of Kaisti 2181 in the presence of algae is observed in our enrichment analysis and our interaction measurement. In contrast, algae inhibits Caulob 3646 in the enrichment analysis (quadrant II in Figure S14) but direct measurements of interactions reveals no effect of algae on its growth.

In contrast, Pseudo 2001, Bacill 530 and Dugane 4601 are part of the abiotic taxa, i.e. they cause motion along PC1 for all communities irrespective of the presence of the algae. One would therefore expect that the presence of algae has no effect on their enrichment dynamics. This is indeed observed when direct interactions are measured for Pseudo 2001 and Bacill 530 (Figure 6A). However, Dugane 4601 does not follow this trend, being strongly suppressed in the presence of algae. Similarly, Sphing 376, which is neither a biotic nor an abiotic taxa, is strongly suppressed in the hybrid communities, whereas the direct measurement of the interaction reveals that it grows better in the presence of algae.

On the whole the interaction measurements are in accord with the sequencing measurements. However, there are exceptions, for which there is not a clear interpretation of the interactions shown in Figure 6A in terms of the location of each isolate on the enrichment plot and their contribution to motion along PC1. The reason for this discrepancy with the co-culture data (Figure 6A) may be that the pairwise interaction is not representative of the emergent dynamics of this OTU and the community context matters qualitatively, or that our isolate is a strain with different ecological interactions than the dominant OTU assigned to that ASV (*35*).

### Bacterial isolates inhibit algal growth

In the previous section, we examined the effect the algae had on the bacteria. Now, using the library of isolates we wanted to understand the effect of bacteria on the growth of *C. reinhardtii*. To study this, we grew *C. reinhardtii* with each isolate individually in conditions identical to those of the enrichment experiment (Methods). We included an algae-only control. Samples were collected from each vial approximately daily and algal abundances were quantified by flow cytometry. Example algal abundance dynamics with and without a bacterial isolate are shown in Figure 6B, C. To quantify the impact on algal growth of a bacterium we computed the area under the growth curve (AUC) (*36*) for algae alone and in the presence of bacteria. If the AUC is lower for algae in the presence of bacteria (AUC^co^) than it is for algae alone (AUC^mono^) this indicates inhibition of algal growth by bacteria via increased lag phase duration, decreased growth rate, or decreased carrying capacity or a combination of any of these. Therefore, to assess the overall impact of bacteria on algal growth we computed the ratio of (AUC^co^)/(AUC^mono^). If this ratio is > 1 it indicates bacterial facilitation of algal growth, if it is < 1, it indicates suppression. Figure 6D shows that only one of the isolates we assayed facilitates algal growth and all others act to inhibit or are neutral. The isolate Ralstonia 4908 has no significant impact on algal growth and we find it is enriched in the presence of algae during the enrichment experiment.

### Bacterial carbon use preferences predict enrichment rates

While our enrichment analysis (Figure 5D) suggests algae predominantly inhibit the biotic bacteria in hybrid communities, other taxa do rise in abundance in the long growth period conditions over 10 rounds of dilution (Figure 2). This suggests that some of these taxa might be consuming algal exudates. A direct measurement of bacterial uptake of algal exudates is beyond the scope of this work, so we decided instead to see if catabolic phenotypes of our isolates were related to their dynamics during enrichment. To investigate this we assayed the growth of each isolate in our library on carbon compounds excreted by *C. reinhardtii*. We determined the compounds excreted by the alga in monoculture using untargeted metabolomics on spent media (Methods). From these data we selected 6 organic carbon compounds that accumulated in time in algal monoculture, and tested growth of all 21 isolates on them and glucose (Figure S16A). To see if these resource preferences had any relation to the dynamics of taxa in our experiment we asked whether or not these resource preferences were statistically related to the enrichment (*r_h,s_*) for each isolate in the library. To do this we performed a regression analysis to predict *r_h,s_* using whether or not each isolate grows on each carbon source as a binary independent variable. This analysis (see SI Section 2.7, Figure S17) showed that for the isolates in the library, *r_h,s_* for *τ*_12_ and *τ*_9_ was predictable (*R^2^* ≈ 0.3 – 0.4) based on carbon resource preferences, but that these preferences had no predictive power for *r_h,s_* measured in the *τ*_3_ or *τ*_6_ hybrid communities (Figure S16). Similarly, the carbon consumption preferences had no predictive power on the enrichment of the isolated taxa in control communities, *r_c,s_* (Figure S16). Carbon sources excreted by the algae had significant coefficients while glucose did not (Figure S17C,D and E). These results suggest that the dynamics of bacteria in the long growth periods are influenced by the catabolic capabilities of each OTU, and that the consumption of algal exudates becomes more important as the frequency of the supply of exogenous nutrients decreases.

## Discussion

Using laboratory enrichment experiments where we varied the frequency of exogenous nutrient supplies and the presence of a phototrophic alga, we demonstrated the relative contributions of abiotic and biotic factors to community assembly. We showed that phototrophs impact community assembly when nutrients are supplied infrequently, suggesting that high levels of allochthonous nutrients should serve to decouple phototrophic microbes from their heterotrophic partners. When initially diverse bacterial consortia from distinct soil samples are grown in long growth periods (9 or 12 days) we find that the presence of the algae drives a convergence in community composition. Thus, phototrophs appear to engineer their local microbiota when exogenous nutrient supplies are diminished. Analysis of changes in relative abundances of bacterial taxa in the presence and absence of the algae shows that this convergence emerges from net algal inhibition of bacterial abundances. Direct measurements of the impact of algae on bacteria confirm this coarse picture, but also show that some bacteria are dependent on algal exudates for growth. Direct measurements also show that the bacterial isolates have a negative impact on algal growth, which we speculate could be a reason for the algae to inhibit taxa.

One striking result of our study is the convergence of bacterial communities in nutrient scarce environments when algae are present. Convergence of community composition despite diverse initial compositions supports the idea that the local environment strongly controls assembly (*5*, *37*). However, this result extends this insight by demonstrating biotic contributions to the environment, namely the presence of specific metabolic phenotypes such as phototrophy, can themselves alter the environment to determine emergent community composition. Our result points to the idea that assembly might be best understood by considering not just abiotic factors but also the traits of those taxa present in the community.

Our analysis of enrichment rates for bacterial taxa with and without algae provides a clear picture of the net impact of the presence of algae on the changes in relative abundances of bacteria. While it is widely appreciated that phototrophs can recruit specific bacterial strains via nutrient exchanges (*8*) our results indicate that negative interactions are important for phototrophic impacts on bacterial community assembly. Overall, our data suggest that algal mediated convergence of bacterial community composition is dominated by net negative impacts on bacterial abundances (Figure 5D). However, our experiments with isolates indicate that some taxa in the community are likely relying on algal exudates to sustain growth (Figure 6D).

The precise interplay between inhibitory interactions and algae recruitment of bacterial taxa via nutrient excretion remains unclear. Our statistical analysis of bacterial catabolic phenotypes and enrichment rates suggests that the ability of bacterial taxa to consume algal exudates is important for determining their dynamics when nutrients are supplied infrequently. However, because our isolate library does not include most of the dominant taxa from different dilution rounds we cannot determine whether those taxa that rise in abundances in later rounds of the long-growth period treatments are doing so because they can exploit autochthonous resources provided by the autotroph or not. Future work should focus on dissecting the interplay between these two processes at a community level.

The mechanisms through which the phototrophs and heterotrophs interact in our experiment are not known. While the regression analysis indicates that consumption of carbon sources secreted by the algae could play a role in selecting taxa, we find that the dominant effect of the algae is to inhibit bacteria to cause community convergence when nutrients are scarce. This suppression could be due to various reasons. One possibility is that acidification of the medium, potentially caused by algae consuming ammonia as a nitrogen source (*38*) (SI section 1.7.1), might inhibit certain taxa. Other reasons could be competition for resources as evidenced by the mutual inhibitory interactions (Figure 6), parasitism (*12*), or oxidative stress (*39*).

Our work provides important context for studies seeking to understand the role of phototrophs (*24*, *29*) and the nutrients (*20*) they supply in structuring complex communities in the wild. Our work suggests that in coastal settings, for example, high levels of allochthonous nutrients might serve to decouple bacteria from the local primary producers. One important caveat to this idea is that it is thought that allochthonous nutrients are more recalcitrant (*21*) than those excreted by autotrophs. As a result, an important extension of the work here is to understand how variation in the identity of the exogenously supplied nutrients impacts the convergence and assembly dynamics observed here. One important limitation of this study is the use of glucose as the exogenously supplied nutrient. It may be that our results depend on the chemical identity of these nutrients, and in particular on which taxa can and cannot consume those nutrients.

More broadly, our study highlights the need for tractable model systems for studying the role of primary producers in structuring microbial consortia. The marine bacterium Prochlorrococcus has served as a powerful model for understanding the ecology of phototrophic microbes in the ocean (*40*). However, Prochlorro-coccus is not to our knowledge genetically tractable and we lack the detailed cell biological insights available in *C. reinhardtii*. In contrast, for *C. reinhardtii*, we lack ecological context and insights (*30*). Our hope is that this study helps take a step towards leveraging the power of model organisms for ecological gain. To that end, utilizing large mutant libraries (*41*) in concert with communities like those assembled here could more deeply connect genomic properties with ecological outcomes.

## Methods

### Enrichment experiments

Soil samples were collected from a restored prairie. The soil was treated with drugs to eliminate eukaryotes and placed in the dark to eliminate native phototrophs. Soil samples from two different locations within the prairie were used to inoculate the experiments. These were designated as Soil A and Soil B. (SI Section 1.1).

A modified version of the freshwater mimicking Taub media (*42*) was used for the experiments at pH 7.2 and with defined carbon, nitrogen and phosphate sources. The components and their concentration are given in Table S1. For the initial growth of *C. reinhardtii* (UTEX 2244 mt+), a modefied version of Tris-Acetate-Phosphate media (TAP) was used. Its preparation, components and concentrations are described in SI Section 1.4.

The enrichment experiment was performed in cylindrical 40 mL vials placed on a 60 position stir plate. An LED light panel was placed between the stir plate and the vials to provide illumination. The LED panel was controlled by a Raspberry Pi via a signal relay and a DC power supply to create light - dark cycles of 12 h duration each. The set-up was in a temperature controlled room, with the temperature maintained at 30 °C (SI Section 1.2).

16s amplicon sequencing of the V4 region was performed on all samples from every other round of serial dilution based on the protocol from the Earth Microbiome Project (EMP) (*43*). All the primers used were obtained from the sequences provided in (*43*). See SI Section 1.8 and (*44*) for more details.

### Principal component analysis on the taxonomic data

PCA was performed on the 16S amplicon sequencing data. However, the date are compositional and need to be analyzed accordingly (*32*, *33*). A standard way to do this is to transform them using a log transformation. But to perform log-transformations, the data cannot have zeros. But sequence data have typically have a large number of zeros. Here we describe how we handle these problems.

#### Aitchison’s transform

Here, we use the center-log-ratio (clr) transformation, suggested by Aitchison (*34*, *45*). If 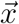 is the vector of the composition of each community, and *x_i_* are the counts of each individual taxa *i*, with *i* ∈ (1, 2,…*N*) and *N* the total number of OTUs found across all communities, then the clr transformation is:

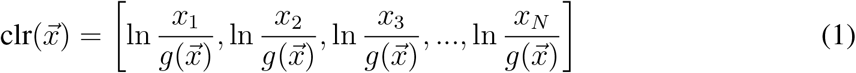

where 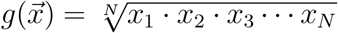 is the geometric mean.

#### Zero removal

From above, it is clear that zero counts will cause the geometric mean to be zero and the transformation will therefore diverge. Although many methods have been suggested to replace zeros (*46*), they are not always suitable for all datasets (*47*). Here, we explain our method of handling the zeros.

First, we add a small number (1) to all counts. It has been shown elsewhere (*48*) that adding small pseudocounts does not affect the downstream analysis process, while effectively removing any zero counts. Then, we clr transform the data, and perform PCA on it. We plot the distribution of the eigenvalues (see Figure S6A). We note that there are 2 eigenvalues that are much higher than the others. Next, we randomly scramble the data, i.e. for each community, we randomly interchange the counts for each taxa (shuffle columns of the data matrix). We clr transform this randomized data and perform PCA on it. This scrambled data does not have any structure, and we expect the high eigenvalues to vanish. The distribution of eigenvalues is shown in Figure S6B. Indeed, we note that the top two eigenvalues disappear.

To remove rare taxa, we plotted the distribution of the frequency of the relative abundances as a function of the maximum relative abundances across all samples at all time points. We observe from Figure S7 that there a large number of taxa that have a low maximum relative abundance across all samples at all times - which means these taxa have close to zero counts in all communities. We set a cutoff at a threshold of −8.2 for the logarithm of the maximum relative abundance, which corresponds to a relative abundance of 0.00027. All taxa that have a logarithm of maximum relative abundance below this number are discarded. With this reduced data, we re-do the above analysis, i.e., we add a small number, 1, to all the counts, take the clr transform and perform PCA. We plot the eigenvalue distribution for this data in Figure S8A. Again, we note that there are only 2 eigenvalues that are much higher than the others. Next, as before, we randomize the data, clr transform it, and perform PCA. The eigenvalue distribution for this dataset shows that both the high eigenvalues vanish as expected (Figure S8B). This disappearance of the two high eigenvalues on randomizing leads us to conclude that the data are well described by 2 eigenmodes. For the rest of the analysis, we use the reduced data, with the extremely rare taxa discarded.

### Experiments with isolates

#### Co-culture experiments via flow cytometry

To study the impact of bacteria on the growth of *C. reinhardtii* we grew *C. reinhardtii* on it’s own, and in co-culture with individual bacterial strains. Bacterial isolates were first grown in R2 media for two days, then diluted into the Taub minimal media. After a further two days of growth, the bacterial isolates were inoculated at a starting OD of 0.0005. Bacterial isolates that did not grow well in liquid media were directly plated and the colonies were used for the experiments. As in the enrichment experiments, *C. reinhardtii* was grown in TAP media and washed three times in the Taub media on saturation. It was then diluted to start the experiments at a low cell density of 100 cells/mL in order to observe the growth curves. Each condition had 3 replicates. The cultures were grown at the same temperature, stirring and light conditions as the enrichment experiment. Samples were collected regularly for measurement of *C. reinhardtii* cell densities. These samples were fixed using the drug cycloheximide (100 *μg* / mL), and stored at 4°C. After a number of samples were collected, they were transferred to a 96 well plate and the cell density was measured using the Attune Nxt 14 flow cytometer housed at the Cytometry and Antibody Technology Core facility at the University of Chicago (Voltages: Forward scatter: 360 V, Side scatter: 200 V, Laser: 260 V). It is a volumetric flow cytometer that can measure flow volume, eliminating the need to use beads for calibration. This resulted in a time series of algal abundances in mono-culture and in co-culture with bacterial isolates. The raw data are shown in Figure S15.

To assess the impact of bacteria on algal growth we computed the area under the abundance verses time curve (AUC) (36) acquired via flow cytometry. The area under the growth curve decreases as either the lag time increase, growth rate decreases, or carrying capacity falls. Therefore, any reduction in the AUC in the presence of bacteria relative to the algae alone indicates an interaction. If the AUC increases this suggests bacterial facilitation of algal growth and if it decreases this suggests inhibition. Therefore, we computed the ratio of the AUC (area with isolate/area without isolate). If this ratio is larger than 1, then that isolate is beneficial for the growth of the algae, and vice versa.

#### Co-culture experiments via plating

To study the effect of *C. reinhardtii* on isolates, each isolate was grown on its own and in the presence of *C. reinhardtii*. Bacterial isolates were first grown in R2 media for two days, then diluted into the Taub minimal media. After a further two days of growth, the bacterial isolates were inoculated at a starting OD of 0.001. Bacterial isolates that did not grow well in liquid media were directly plated and the colonies were used for the experiments. As in the enrichment experiments, *C. reinhardtii* was grown in TAP media and washed three times in the Taub media on saturation. It was then diluted to start the experiments at a cell density of 10^5^ cells/mL. After 14 days of growth in the same conditions as the enrichment experiment, the communities were serially diluted and 10 *μ*L of the serially diluted cultures were plated on agar plates (*49*) to measure the end point cell density of the isolates when grown on their own and when grown in the presence of algae. The lowest dilution factor used was 10^-4^. Due to this, the detection limit of the cell density in this method is 1 × 10^6^cells/mL.

#### Chemical analysis of algal exudates

To find the organic carbon compounds excreted by *C. reinhardtii*, we grew the algae axenically in three replicates (without any soil communities, or other bacteria). Samples were collected at the end of 3 days, 6 days, 9 days and 12 days, corresponding to the growth periods used in the experiment. The samples were then sent for analysis using GC-MS to the Roy Carver biotechnology center at the University of Illinois at Urbana Champaign (Agilent 7890A GC/5975C MS). The resulting data indicated the presence of about 40 different organic compounds (Dataset7). To infer which of these compounds were being secreted increasingly with time, we subtracted the baseline compounds found in the media, and then plotted the measurements as a function of time. We then performed a linear regression, and found which compounds had a significant positive slope. The data from GC-MS is provided in the Dataset 7 and the compounds that are secreted in significant amounts are listed in Table S7.

## Supporting information

Dataset_1_16s_counts_data

Dataset_2_16s_phylogeny

Dataset_3_OTU_counts_data

Dataset_4_OTU_phylogeny

Dataset_5_p_values_PC1_movement

Dataset_6_p_values_PC1_movement_no_thresholding

Dataset_7_GCMS

Dataset_8_taxonomy_isolates

Dataser_9_treefile_rare_taxa_removed

## Acknowledgments

SK would like to Acknowledge funding from the National Science Foundation (BIO 2117477). KHP would like to acknowledge funding from the Center for Physics of Living Cells, UIUC. The authors thank Derek Ping for help with the experiments, University of Chicago’s Cytometry and Antibody Technology (CAT) facility for flow cytometry measurements, the Raymond J. Carver Biotechnology Center at the University of Illinois at Urbana-Champaign for GC-MS measurements, and Karna Gowda for useful comments and discussion.

