## Supplementary Appendix for "Algae drive convergent bacterial community assembly when nutrients are scarce"

### 1 Experimental methods

#### 1.1 Collecting and processing soil samples

Soil samples were collected from a restored prairie (Meadowbrook Park, Urbana, IL) located at 40˚04’42.9”N and 88˚12’22.3”W. Sterile scoopulas were used to dig holes about 5 cm deep and soil from the bottom of the holes was collected into separate sterile Falcon tubes for each hole. The samples were then taken to the lab. About 5 g of each soil type was weighed into fresh Falcon tubes. 10 mL double distilled water was added to each Falcon tube. The soil was soft and broke down into fine particles on vortex mixing. This soil suspension was then stored at 4°C.

To extract bacterial communities from the soil samples, the Falcon tube was vortexed thoroughly and the large soil particles were allowed to settle. However, very small soil particles and bacteria remain suspended in the supernatant. This supernatant containing bacteria and small soil particles was collected and centrifuged at 7000 rpm for 5 min. The pellets contained bacteria and small soil particles. The supernatant was discarded and the soil bacterial pellet was re-suspended in media without organic carbon. This was then distributed into sterile test tubes so that each test tube received no more than 2.5 mL total. To each test tube, 200 $\mu$g/mL of the drug cycloheximide was added to kill any eukaryotes. Further, 0.02 mg/mL of the fungicide Nystatin was added to kill any fungi. The test tubes were then wrapped in Aluminum foil to prevent light from entering the soil sample, thus prohibiting the growth of any native phototrophs present in the community. These wrapped test tubes were then placed in a shaker-incubator, where they were shaken at 225 rpm and incubated at 30°C for 2 days.

After two days of incubation, the samples were collected from the test tubes and centrifuged at 7000 rpm for 5 min, causing the bacteria and remaining small soil particles to form a pellet. The supernatant, which contained the drugs, was discarded. The bacterial community was then re-suspended in an equal amount of fresh media without organic carbon. The samples were then ready for further use. The communities at this stage will be referred to as initial soil inoculates.

#### 1.2 Experimental setup for enrichment experiments

A multi-position stir plate (2mag-USA) was used to to stir 60 vials simultaneously. A multi position vial holder was custom designed to hold 60 vials each of 40 mL volume. An LED light panel (Good Earth Lighting, Item: 805392, Model: LF1089-PEW-28LF4-G) was placed between the vial holder and the multi-position stir plate to provide uniform illumination from below. The light panel was powered by a DC power supply, via a signal relay mechanism. The signal relay was in turn controlled by a Raspberry Pi, which allowed us to turn the lights on and off every 12 h, thus simulating day - night cycles. The vial holder was enclosed in a thick black cardboard box to ensure that external light changes did not affect the experiment. The entire set up was placed in an incubated hot room set to 30°C.

The input voltage (19 V) and current (1 A) to the LED light sheet was set on the DC power supply so that the average light intensity at each position was 1050 lux with a standard deviation of 50 lux across all positions as measured using a luxmeter. This is approximately 30 μmol/$m^2$/s as measured by an LI-COR light meter. The color temperature of the LED light panel was 3975K (bright white).

The vials are made of glass, are cylindrical and have a volume of 40 mL. Each vial contained a stir bar. In the experiments, 20 mL of cell culture was used. The vials were closed using sterile foam stoppers, which permit airflow but maintained sterility. We used large vials, rather than microtiter plates, due to the long timescales of the growth periods where evaporation would cause excessive water loss in smaller volume cultures.

#### 1.3 Measurement of evaporation

Evaporation was measured by measuring weight of two vials with water every 24 h in the experimental conditions, i.e, with the foam stopper, with stirring, and with the LED lights on. Subtracting the initial weight of the vial, and using the density of water, the volume of water evaporated was calculated to be about 0.38 mL/day. This means that for the longest growth period of 12 days, the total volume of water evaporated would be 4.5 mL.

#### 1.4 Tris Acetate Phosphate (TAP) media

TAP media was used to start the *C. reinhardtii* cultures from the frozen stocks. First, a 2X Filner Beijernicks solution is made by dissolving 8 g $NH_4Cl$, 1 g $CaCl_2.2H_2O$ and 2 g $MgSO_4.7H_2O$ in 500 mL MilliQ water. This stock solution is then autoclaved and stored at 4°C.

Next, a stock of trace mineral solution is prepared. We start with dissolving 5g of disodium EDTA in 400 mL double distilled water by heating and constant stirring. The pH is set to 6.5 using 5N NaOH. Then, the following compounds are added in the given order, ensuring that each compound dissolves before adding the next: $FeSO_4.7H_2O$ 0.5 g, $ZnSO_4.7H_2O$ 2.2 g, $H_3BO_3$ 1.14 g. $MnCl_2.4H_2O$ 0.51 g, $CuSO_4.5H_2O$ 0.016 g, $Na_2MoO_4.2H_2O$ 0.073 g, and $Co(NO_3)_2.6H_2O$ 0.0196 g. The volume is then made up to 500 mL and autoclaved.

Next, a phosphate stock is prepared by adding 6.8 g of $KH_2PO_4$ and 8.7 g of $K_2HPO_4$ to 50 ml MilliQ water and autoclaving.

Finally, TAP media is prepared by combining 12.5 mL of the 2X Filner's Beijernick's solution, 0.5 mL of the phosphate stock, 2.5 mL of the trace mineral solution, 1.21 g of Tris- base, and 0.5 mL of glacial acetic acid. The volume is brought to 500 mL, the pH adjusted to 7.2, and the solution is autoclaved. It is stored at room temperature.

#### 1.5 Starting the enrichment experiment

A lab strain of *C. reinhardtii*, UTEX2244 mt+, was grown from stock in TAP media for three days in continuous light conditions, with shaking, and at 30 °C. After three days' growth, the culture was centrifuged at 5000 rpm for two minutes. The supernatant was discarded and the cells re-suspended in the Taub media. They were then centrifuged again, at the 5000 rpm for 2 min. Again, the supernatant was replaced with fresh Taub media. This process was repeated again 3 times in total.

The cells were then counted using a hemocytometer and the appropriate dilution to get $1 \times 10^5$ cells/mL initial cell density was computed. 250 $\mu$L of the initial soil inoculates, which are the source of heterotrophs for our experiment, was added to each vial. The remaining culture volume (total volume 20 mL) was comprised of the Taub medium described in Methods. Once the vials were assembled, they were placed in the multi-position vial holder and the light cycle was started via the DC power supply controlled by the signal relay in turn controlled by a Raspberry Pi.

#### 1.6 Propagating the experiment

After the vials have grown for the number of days specified by their growth period, a 1:64 dilution was performed into fresh Taub media. This process was repeated until communities of all growth periods have completed 10 rounds of serial dilution. At the end of each serial dilution, samples were collected to measure Chlorophyll fluorescence and optical density. Samples were also preserved with 25% glycerol at −80 °C and for 16s sequencing at −20 °C. Note that the $\tau_{12}$ and $\tau_9$ control communities had 1 and 2 replicates contaminated by *C. reinhardtii* respectively, and these contaminated samples are not used in the analysis.

#### 1.7 Optical Density and Chlorophyll fluorescence measurements

At the end of each round of serial dilution, samples from the communities were transferred to the wells of a 48 well plate. The optical density (OD) was determined by measuring absorbance at 600 nm of the samples in the 48 well plate using a plate reader (BMG Labtech, Clariostar). The OD600 of the Taub media was measured separately, and used as background for background subtraction. The background subtracted OD600 measurements for the hybrid communities are shown in Figure S1A. We note that the OD600 of the $\tau_{12}$ hybrid communities is much higher than that for the $\tau_3$ and $\tau_6$ hybrid communities by round 10 in both soil samples. In soil A, the $\tau_9$ hybrid communities also have a higher OD600 by round 10 compared to the $\tau_3$ and $\tau_6$ hybrid communities. In comparison, Figure S1B shows the background subtracted OD600 measurements for the control communities. By dilution round 10, control communities in both the soil samples reach a similar OD600 of about 0.3 irrespective of the growth period, despite starting at different OD600 values initially.

Auto-fluorescence of chlorophyll was used to quantify chlorophyll content by measuring fluorescence in a Clariostar Plate Reader at excitation wavelength of $482 \pm 20$ nm, emission wavelength of $690 \pm 20$ nm using a dichroic at 585 nm. The results of the measurements for hybrid communities are shown in Figure S1C. In Soil A, the $\tau_{12}$ communities have a higher chlorophyll content. However no other trend is visible.

In order to compare the OD600 of the hybrid communities without the contribution of *C. reinhardtii*, we plotted the ratio of OD600:Chlorophyll fluorescence in Figure S1D. This shows that $\tau_3$ hybrid communities have a higher OD:Chlorophyll content compared to hybrid communities of other growth periods for both soil samples. In other words, for the $\tau_{12}$ hybrid communities, which had higher OD600 values, the high OD is a result of the *C. reinhardtii* abundance. Further, the hybrid communities with the shorter growth periods have more bacteria per chlorophyll content than the hybrid communities with the longer growth period. This is consistent with the loss in diversity observed (Figure 3) in the hybrid communities with longer growth period.

### 1.7.1 pH

The approximate pH of $\tau_{12}$ Soil B communities was measured at the end of round 7 of serial dilution using pH paper. We found that the pH of the hybrid communities had dropped to between 5 and 6, whereas the pH of the control communities remained at around 6.4.

Since the pH of hybrid communities dropped a lot more, we hypothesized that *C. reinhardtii* could be responsible. In a media with $NH_4Cl$, algal growth acidifies the media by taking up $NH_3$ and leaving behind $H^+$ ions [1]. To test this hypothesis, we grew *C. reinhardtii* in media with two different $NH_4Cl$ concentrations. One had the concentration used in the experiment (8 mM) while the other had a lower concentration of $NH_4Cl$ of 2 mM. In both media, the initial pH was set to 7.2 The pH of the culture was measured every 3 days for 12 days. The drop in pH was steeper in the media with 8 mM $NH_4Cl$ than in the media where the concentration was 2 mM. Further by the end of 12 days, the pH had dropped to 5.7 in the media with 8 mM $NH_4Cl$ and seemed to be declining with time (negative slope of pH vs time), whereas in the media with 2 mM $NH_4Cl$, the pH had dropped to 6 and seemed to stabilize there (slope of pH vs time was about zero).

This indicated that *C. reinhardtti* causes the drop in pH due to the consumption of $NH_4Cl$. The sharper drop in pH seen in the hybrid communities could be due to the contribution of the heterotrophs to the acidification process.

#### 1.8 16s Sequencing

##### 1.8.1 DNA extraction

DNA was extracted using Qiagen's DNeasy 96 well Blood and Tissue kit. A modified version of the protocol was used. Just prior to starting the extraction process, a lysis buffer (see Table S2) was freshly prepared. The samples frozen at -20°C were thawed and centrifuged for 15 min at 21000 rcf. The supernatant was discarded and the pellets were re-suspended in 180 $\mu$L lysis buffer. The samples were then incubated for 30 min at 37°C. 25 $\mu$L proteinase K and of 200 $\mu$L Buffer AL (without ethanol) were added to the samples. The samples were then incubated at 56°C for 1 h. Next, 200 $\mu$L ethanol was added to the samples and mixed by vortexing. The samples were then centrifuged briefly (allowed to reach 3000 rpm and then stopped) to collect any solution accumulated in the caps.

A DNeasy 96 well plate was placed on top of an S block (both provided with the kit). The lysate was transferred to the wells of the 96 well plate. The plate was then sealed with an AirPore Tape sheet and centrifuged at 4000 rpm for 15 min. The tape sheet was then removed and 500 $\mu$L buffer AW1 was added to all the wells. The plate was then sealed again with a new tape sheet and centrifuged for 10 min at 4000 rpm. After centrifugation, the tape was removed and 500 $\mu$L buffer AW2 was added to all samples. The plate was centrifuged for 20 min at 4000 rpm without a tape sheet.

The DNeasy 96 well plate was then placed on a rack of elution microtubes, and 100 $\mu$L of elution buffer AE added to all samples. The samples were incubated for 1 min at room temperature and centrifuged at 4000 rpm for 3 min. This step is repeated again. The extracted DNA is in the elution microtube, which is then covered and stored for future use.

##### 1.8.2 DNA extraction from soil

The kit used above to extract DNA from liquid cultures was not suitable for extracting DNA from bacteria present in soil. So we used Qiagen's DNeasy Power Soil Pro kit to extract DNA from the initial soil inoculates (the communities after being extracted from soil and after treatment with drugs for 2 days in the dark) that were used to initiate the experiments, preserved after the treatment with drugs. The beads of the PowerBead Pro tube were removed and the thawed soil sample (preserved in liquid, about 20 mg) was added to the tubes. The tubes were centrifuged at 1000 rcf for 30 s, the supernatant was removed. The beads were then added back into the tube. 800 $\mu$L of solution CD1 was added and the tube was vortexed. The tubes were then fixed horizontally into a vortex adaptor and vortexed for 10 min to homogenize the samples. The tubes were then centrifuged at 15000 rcf for 1 min. The supernatant was transferred into micro-centrifuge tubes. 200 $\mu$L of solution CD2 was added and the tubes were vortexed for 5 s. Then the tubes were centrifuged at 15000 rcf for 1 min. The supernatant was transferred into a clean micro-centrifuge tube. 600 $\mu$L of solution CD3 was then added and the tubes vortexed.

650 $\mu$L of the lysate was loaded onto an MB spin column and centrifuges at 15000 rcf for 1 min. The flow through was discarded, and the process was repeated until all the lysate was passed through the column. The MB spin column was then placed in a 2 mL collection tube and 500 $\mu$L of solution EA was added to it. After centrifugation at 15000 rcf for 1 min, the flow through was discarded and the spin column placed into the same collection tube. 500 $\mu$L of Solution C5 was then added and centrifuged at 15000 rcf for 1 min. The flow through was again discarded the spin column was placed in a fresh collection tube. The tube was then centrifuged at 16000 rcf for 2 min and the spin column was placed in an elution Tube. 75 $\mu$L of the Solution C6 was added to the center of the filter membrane and the tube was centrifuged at 15000 rcf for 1 min. The flow through contained the DNA and was stored at -20°C.

##### 1.8.3 Library prep

Library prep was performed according to the protocol suggested by Illumina. In particular, first, to ensure that the DNA extraction worked, the DNA concentration was quantified in all samples using Invitrogen's Qubit dsDNA BR Assay kit, adapted to a 96 well plate version.

**Qubit**

Enough working solution was prepared by mixing Qubit dsDNA BR buffer with Qubit dsDNA BR reagent in a 1:200 ratio. Using the provided standard solutions, one 100 ng/$\mu$L and the other 0ng/$\mu$L, serial dilutions were performed to obtain a range of known DNA concentrations. 195 $\mu$L of the working solution, and 5 $\mu$L of the known DNA concentrations were added to the wells of the 96 well plate. The plate was vortxed briefly, allowed to incubate, and placed in a plate reader. The excitation wavelength is 485 nm and the emission wavelength is 530 nm. The wells were scanned using these parameters and calibration curve was obtained.

Once the calibration curve was obtained, the the same procedure described above was used to measure the fluorescence of the unknown samples, using the same working solution. From the calibration curves, the DNA concentration in the samples was inferred.

**PCR**

Once it was confirmed that the DNA extraction process had worked, PCR was performed on the extracted DNA in accordance with the Earth Microbiome protocol [2]. All samples received the same forward primer but unique reverse primers. Each reverse primer has a unique barcode, which can be later used to demultiplex reads belonging to each community. The reaction were performed in triplicate (see Table S3 for the volumes) with a total volume of 25 $\mu$L each. The master mix used was the Platinum Hot Start Master mix. See Table S4 for thermocycler settings.

After PCR was performed on all samples, the triplicates were pooled. The DNA concentration of each sample was measured using the Qubit dsDNA BR Assay kit as described above. For each sample, the volume required to obtain 240 ng of DNA was calculated. This volume was then collected from the respective samples and pooled. The total volume of the pool was also calculated.

The pooled PCR products were then cleaned using QIAquick PCR purification kit. Ethanol was added to buffer PE prior to use. pH indicator was added to the buffer PB in a ratio of 1:250. Yellow color of the resulting mixture indicated a pH $> 7.5$. 5 volumes of the buffer PB were added to the 1 volume of the PCR products and mixed. A QIAquick spin column was placed in a collection tube. The sample was then added to the spin column and centrifuged at 17900 rcf for 45 s. The flow through was discarded and the spin column was placed back in the same collection tube. 750 $\mu$L of the buffer PE was added to the spin column and centrifuged at 17900 rcf for 45 s. The flowthrough was discarded and the spin column was placed back in the same tube and centrifuged at 17900 rcf for 1 min. Then the spin column was placed in a clean microcentrifuge tube. 30 $\mu$L of the buffer EB was added to the center of the spin column's membrane and let stand for a minute. The column was then centrifuged at 17900 rcf for 1 min. The flow through contained the cleaned and pooled PCR products.

The final concentration of the cleaned and pooled PCR products was measured using the Qubit dsDNA BR Assay kit as described above. The concentration of the DNA was estimated using the following formula [3]:

$$\frac{\text{concentration in ng}/\mu L}{660\text{ g/mol} \times \text{average library size}} \times 10^6 = \text{concentration in nM} \tag{S1}$$

For our experiments, the average library size was 390. Using resuspension buffer, the library was diluted to a concentration of 4 nM.

##### 1.8.4 Sequencing

A MiSeq Illumina machine was used with a v2 300 cycle kit. The sequencing was carried out in accordance with [3]. Fresh 0.2 N NaOH was prepared and used to dilute the 4 nM pooled library to 2 nM. It was then vortexed and incubated at room temperature to 5 min to allow denaturation. The denatured DNA was then diluted to 20 pM using pre-chilled HT1 buffer and then further to 8 pM using the same pre-chilled HT1 buffer.

Next, a PhiX control was diluted to 4 nM using resuspension buffer. Using the freshly prepared NaOH, PhiX was diluted down to 2 nM, which was then vortexed and incubated at room temperature to allow denaturation. The, using pre-chilled HT1 buffer, the library is diluted to a 20 pM library, and them further to 8 pM.

Finally, the denatured PhiX and the denatured sample library are pooled so that PhiX makes up 5% of the mixture. The mixture was then placed in a heatblock at 96°C for 2 min.

A .csv file was prepared with the information about the samples and the kit details. This file was loaded onto the MiSeq machine. The thawed cartridge was flipped up and down a few times. The foil marked for sample was pierced using a pipette tip and the sample loaded into it. Reservoirs 12, 13 and 14 were also pierced with clean pipette tips. 3.4 $\mu$L of 100 $\mu$M Index sequencing primer was added to reservoir 13, 3.4 $\mu$L of read 1 sequencing primer was added to reservoir 12, and 3.4 $\mu$L of read 2 sequencing primer was added to reservoir 14. The kit was then loaded into the machine, and the run was started.

#### 1.9 Isolation of strains

To isolate individual strains from the communities, the stock preserved in glycerol was used. These stocks were grown in vials in Taub media in the presence of *C. reinhardtii* at the same concentration as in the enrichment experiments. The communities were allowed to grow for the same number of days as in the experiment with cycles of light and dark i.e., a 3 day community was re-grown for 3 days, a 9 day community for 9 days and so on. At the end of the growth, the communities were diluted and plated on R2 agar plates and Taub agar plates.

When individual colonies grew, each distinct looking colony was selected and grown on a fresh plate. This process was repeated until it was clear visually that the the strains were pure.

Colony PCR (see Table S5 for PCR conditions and Table S6 for thermocycler settings) was performed on these strains by picking a single colony and dissolving it in 100 $\mu$L PCR grade water. The same forward primer and one of the reverse primers used in the EMP protocol were used. The master mix used for the EMP protocol was also used here.The PCR products were sent for Sanger sequencing. The results of the Sanger sequencing were compared with the results of Illumina MiSeq sequencing to assign the right label to each isolated strain. To do this, we used Biopython [4] SeqIO's "abi-trim" option, which trimmed the input Sagner sequence based on the quality scores using Mott's algorithm. To align the Sanger sequence with the 16s MiSeq reads, we used Biopython's local alignment, with a match score being the score of identical characters, and with penalties for opening and extending gaps. In particular, identical characters were given a score of 4, 2 points were deducted for every non-identical character, 2 points were deducted for opening a gap and 1 point was deduced for extending an open gap.

Through this process, 21 unique bacterial isolates were obtained. See Dataset 8 for more details of the isolated taxa.

##### 1.10 Carbon utilization assays

Individual isolates were grown in 2 mL R2 media in test tubes at 30 °C with constant shaking for 48h. After 48 h, 1 mL of the cell cultures were centrifuged at 21300 rcf for 2 min. The cells were washed three times in the Taub media. Then, the optical density of cell cultures (absorbance at 600 nm) was measured. The volume required to have a starting OD of 0.01 in a 750 $\mu$L solution was calculated for each isolate to get three replicate communities. In a 48 well plate, the calculated volumes of the cell cultures were added into 3 wells per isolate. The rest of the 750 $\mu$L volume was made up by the Taub media. The plate was then sealed with parafilm and the placed in the plate reader with incubation at 30°C. Absorbance at 600 nm was then measured continuously every 10 min for about 4 days. In between the measurements, the plate underwent orbital shaking at 400 rpm.

Seven different sources of organic carbon were used in the Taub media. One was glucose, and the other 6 were chosen from the GC-MS data of the algal spent media (Methods). The growth curves of isolates on all 7 of these carbon sources were measured using the method described above.

The growth rate and maximum OD reached were calculated from the growth curves by plotting the data on a semi-log scale, after background subtraction. A small number was added to all the data to make all the points positive. The curves were then fit using a spline. The maximum yield (maximum OD600) was found, and the maximum growth rate was inferred from the first derivative of the spline fit.

To binarize the data for the LASSO regression analysis (Section 2.7), any isolate with a maximum OD600 of 0.05 or higher on a particular carbon source was considered to have utilized that carbon source. If the isolate did not reach an OD600 of 0.05, it was considered to be incapable of growing on that carbon source.

#### 2 Data analysis

##### 2.1 16s sequence data processing

Fastq files were generated on Illumina's MiSeq sequencing machine after the run was completed. This gave 3 files, one each for the forward, reverse and barcode reads.The fastq files were then analyzed downstream using Qiime2 [5] and DADA2 [6]. The fastq files are de-multiplexed in Qiime2 using a file assigning barcodes to sample names. The reads were then passed through a quality control check and phiX reads were filtered.

##### 2.1.1 Amplicon Sequence Variants (ASVs)

In order to obtain ASVs, the de-multiplexed reads were analyzed using DADA2 in R using the DADA2 pipeline [7]. The quality of the reads, both forward and reverse, were plotted and it was found that due to the good quality of reads, trimming was not necessary. The reads were then filtered for the number of expected errors. The error rates were learnt from the dataset for the forward and reverse reads. Next, dereplication was performed on the forward and reverse reads to combine the identical reads into unique sequences, with counts information. Using the error rates calculated above, the number of unique sequences were inferred from the dereplicated reads. Next, the forward and reverse reads were merged, a sequence table constructed, and chimeras removed. Finally, taxonomy was assigned by using SILVA's reference database (v128). The ASV level composition of the communities is shown in Figure S3. See Dataset 1 for the counts data and Dataset 2 for the phylogeny data at the ASV level.

##### 2.1.2 Operational Taxonomic Units (OTUs)

In order to obtain OTUs, the demultiplexed files were denoised in Qiime2 using the DADA2 plugin [8]. Then, using Qiime2's vsearch option, the reads were clustered at 97.5% similarity to obtain the count table for the OTUs. This counts table was exported for further analysis. The taxonomy was assigned using Greengenes database on Qiime2. See Dataset 3 for the counts data and Dataset 4 for the phylogeny data at the OTU level. The fasta file for these data can be freely accessed at https://doi.org/10.5281/zenodo.6760409.

#### 2.2 Shannon diversity

Shannon diversity was calculated for each community on all the data i.e., without removing rare taxa. This diversity index is calculated as:

$$H = -\sum_{i} p_i log_e p_i \tag{S2}$$

Here H is the Shannon diversity index of a community, $p_i$ is the probability of finding taxa $i$ in a given community (relative abundance), and the summation is over all the taxa.

#### 2.3 Distance metrics

The counts per taxa was converted to relative abundance for each community by dividing the number of counts for each taxa by the total number of counts for that community. To compare the similarities and differences between the communities, several metrics were used. They are described here.

##### 2.3.1 Jensen-Shannon divergence

Given that the data are in the form of relative abundances, each community can be considered as a normalized probability distribution of the taxa. In this case, Jensen-Shannon divergence (JSD) is a good metric to be used [9], because it is bounded, has the capability to be weighted and is symmetric [9]. The JSD between two normalized probability distributions, $X$ and $Y$, is given by Equation S3, where H is the Shannon entropy of the probability distribution and $\pi_1$ and $\pi_2$ are the weights for the two distributions.

$$J_{X,Y} = H(\pi_1 X + \pi_2 Y) - \pi_1 H(X) - \pi_2 H(Y) \tag{S3}$$

If the distribution $X$ is given by $X = \{x_i\}$, where $x_i$ represent the normalized probability of finding the value $x_i$ in the probability distribution $X$, the entropy $H$ of the probability distribution $X$ is defined by Equation S4.

$$H(X) = -\sum x_i \log x_i \tag{S4}$$

Here, we set the weights to be $\pi_1 = \pi_2 = \frac{1}{2}$. Using Equation S3, we can now define the Jensen Shannon divergence between the relative abundance composition of the different communities.

In order to assess if the hybrid communities are indeed different from control communities, we found the JSD between all pairs of hybrid and control communities. If $H_{i,g}^{r,s}$ is the $i^{th}$ replicate of a hybrid community of soil type $s$, with growth period $g$ and at dilution round $r$, and $C_{j,g}^{r,s}$ is the $j^{th}$ replicate of a control community of soil type $s$, with growth period $g$ and at dilution round $r$, we first found the set of distances $H_{i,g}^{r.s}$-$C_{j,g}^{r,s}$, for all pairs $i, j$. We term this the inter community distances. Next, as a comparison, we found the distances within each hybrid community of a given growth period and dilution round, i.e. we found the set of distances $H_{i,g}^{r,s}$-$H_{j,g}^{r,s}$ where $i, j \in \{1, 2, 3, 4, 5\}$ and $i \neq j$. We term these the intra hybrid community distances. We now have a distribution of inter community distances and intra hybrid community distances. We used bootstrapping (sampling with replacement) to generate different instances of these two distributions. At each instance, we find the difference in medians of the two distributions, $d$. So, at the end, we have a distribution of difference in these medians. Figure S5 shows the distribution of these difference in medians for the each soil sample, growth period, and dilution round. We observe that $d$ is larger than zero (p-value $< 0.002$, with the null hypothesis that $d$ is equal to or less than zero) for the $\tau_{12}$ and $\tau_9$ communities in both soil samples by dilution round 10, whereas for $\tau_3$ communities $d$ is close to zero even by dilution round 10 (p-value 0.13). This means that the hybrid communities are much more distinct from the control communities for the 12 and 9 day growth periods than the 3 day growth periods. This shows that the impact of *C. reinhardtii* on community assembly is stronger for the longer growth periods where the supply of external nutrients is infrequent.

###### 2.3.2 Aitchison's distance metric

Aitchison's distance metric is a way to measure taxonomic distances between communities when the data are compositional, and has some advantages over Bray - Curtis and Jensen Shannon distance metric that are discussed elsewhere[10].

The data are first clr transformed, and then the Euclidean distance is found between all community pairs. The distances are then embedded on 2 dimensions using Multi-dimensional embedding (Figure S9 top panel). The stress of the multi-dimensional embedding is shown in the bottom panel of Figure S9.

The embedding of the Aitchison distance shows similar results as the PCA analysis, with the hybrid communities of the longer growth periods converging to become taxonomically similar, while the other communities do not.

###### 2.3.3 Unifrac

The distance measures in PCA or the Aitchison analysis above do not account for the phylogenetics of the taxa. We wanted to check whether our conclusions were robust to including phylogenetic relationships. To account for the phylogenetic information in finding the distances, we used the unweighted Unifrac distance metric. For this, a tree file was first created by uploading the fasta file of the sequences to SILVA's Alignment, Classification and Tree Service [11]. The tree, along with the counts matrix was then used as input for R's phyloseq package to create a phyloseq object. The newick treefile is available as Dataset 9. Using this object, the unweighted Unifrac distances were computed on R [12] between all pairs of samples. These distances were then embedded in 2 dimensions using multi-dimensional scaling. The results are shown in panel A of Figure S10 and the stress of the embedding is shown in panel B.

The embedding of the Unifrac distance shows similar results as the PCA analysis, with the hybrid communities of the longer growth periods converging to become taxonomically similar, while the other communities do not.

#### 2.4 PCA without removing rare taxa

For completeness, we performed PCA on the data without removing rare taxa. The results are shown in Figure S6C, plotted similar to Figure 6. As in the case where the rare taxa were removed, $\tau_9$ and $\tau_{12}$ hybrid communities converge, with communities of Soil sample B moving downwards along PC2. The control communities and the $\tau_3$ and $\tau_6$ communities do not converge. All the communities, irrespective of growth period and presence of algae move along PC1.

#### 2.5 Displacement along PC2

In order to find the taxa responsible for the displacement of the $\tau_{12}$ and $\tau_9$ hybrid communities of soil sample B, we first note that the eigenvector along PC2 is a vector of loadings along PC2 corresponding to each taxon $k$, i.e., $\vec{PC}2 = \{p_1^2, p_2^2, p_3^2, ..., p_k^2, ...\}$, where $p_k^2$ is the loading corresponding to taxon $k$ along PC2 (denoted by the superscript 2). The composition of a community can also be represented as a vector of the abundances of the taxa in it, i.e., $\vec{c} = \{x^1, x^2, x^3, ..., x^k, ...\}$, where $\vec{c}$ represents a community and $x^k$ is the mean subtracted, clr transformed abundance corresponding to taxon $k$. The displacement of a community along PC2 is actually a summation over all the taxa, i.e. $d_\tau = |\langle \vec{c}^{\tau}_{B,2} | \vec{PC}2 \rangle - \langle \vec{c}^{\tau}_{B,10} | \vec{PC}2 \rangle| = \sum_k |(x^{k,\tau}_{B,2} - x^{k,\tau}_{B,10})| p_k^2$, where $\vec{c}^{\tau}_{B,r}$ represents the hybrid community at round r, soil sample B, with growth period $\tau$ (here, either 9 or 12 days). Here, the notation $\langle \vec{p} | \vec{q} \rangle$ denotes the inner product of the two vectors $\vec{p}$ and $\vec{q}$ . The summation is over all the taxa $k$ and $p_k^2$ is the eigenvector loading corresponding to taxon $k$ along $\vec{PC}2$, and $x^{k,\tau}_{B,r}$ is the mean subtracted, clr transformed abundance corresponding to taxon $k$ at round $r$, of soil sample B, with growth period $\tau$. With this we can find the contribution of each taxa to the displacement of the corresponding communities. The contribution of taxa $k$ to the displacement of a community of soil sample B along PC2 is $o_k = |(x^{k,\tau}_{B,2} - x^{k,\tau}_{B,10})| p_k^2$.

We computed $o_k$ for all taxa $k$ for each replicate community. The median across the replicates was used as the contribution of the taxa to displacement along PC2 for that community. The cumulative distribution was obtained for all taxa for each growth period separately, and a cut-off value was chosen to be at 97.5 percentile. The taxa that had a $o_k$ higher than this cutoff were retained as biotic taxa.

We performed this analysis on the dataset with all the taxa (Figure S6D) and on the dataset after removing rare taxa, obtaining similar results.

#### 2.6 Displacement along PC1

The contribution of each taxa to the displacement along PC1 is found in a similar manner. As before, the eigenvector along PC1 is a vector of loadings along PC1 corresponding to each taxon $k$, i.e., $\vec{PC}1 = \{p_1^1, p_2^1, p_3^1, ..., p_k^1, ...\}$, where $p_k^1$ is the loading corresponding to taxon $k$ along PC1 (denoted by the superscript 1). The composition of a community can also be represented as a vector of the abundances of the taxa in it, i.e., $\vec{c} = \{x^1, x^2, x^3, ..., x^k, ...\}$, where $\vec{c}$ represents a community and $x^k$ is the mean subtracted, clr transformed abundance corresponding to taxon $k$. Here, we find the contribution of the taxa to the displacement along PC1 for all 4 growth periods, for both hybrid and control communities. The displacement along PC1 is given by $d_\tau = |\langle \vec{c}^{\tau}_{s,2} | \vec{PC1} \rangle - \langle \vec{c}^{\tau}_{s,10} | \vec{PC1} \rangle| = \sum_k |(x^{k,\tau}_{s,2} - x^{k,\tau}_{s,10})| p^1_k$, where $\vec{c}^{\tau}_{s,r}$ represents the community at round r, soil sample $s$, with growth period $\tau$. The summation is over all the taxa $k$ and $p^1_k$ is the eigenvector loading corresponding to taxon $k$ along PC1, and $x^{k,\tau}_{s,r}$ is the mean subtracted, clr transformed abundance data corresponding to taxa $k$ at round $r$, of soil sample $s$, with growth period $\tau$.

With this we can find the contribution of each taxa to the displacement of the corresponding communities. The contribution of taxon $k$ to the displacement of a community of soil sample $s$, growth period $\tau$ along PC1 is $o_k = |(x^{k,\tau}_{s,2} - x^{k,\tau}_{s,10})| p^1_k$.

Using the eignevector loadings along PC1, we find the $c_k$ for all taxa $k$ for each replicate community. The median value of $o_k$ across the replicates was assigned as the contribution of that taxa for that community's displacement along PC1. The cumulative distribution of the contributions of all taxa for each growth period was obtained and a cut-off value chosen to be at 97.5% percentile. The taxa that have a $c_k$ higher than this cutoff were retained as abiotic taxa.

In order to test whether the abiotic taxa selected depend on the presence or absence of *C. reinhardtii*, we performed Fisher's exact test [13] on the selected taxa. The null hypothesis was that the selected taxa are equally likely to belong to the hybrid and the control communities. We found the p-value for each selected OTU. There were 77 OTUs selected in total for the thresholded data. To account for the multiple testing, we used the Bonferroni correction [14] to obtain a corrected cutoff for the p-value to be 0.000649. None of the selected taxa had a p-value lower than this in either soil sample (Dataset 5). This implied that the null hypothesis could not be refuted, which means that the selected taxa had an equal probability to be selected from either the hybrid or the control communities. This result means that those taxa that contribute to motion along PC1 are not specific to hybrid or control communities and we therefore refer to them as abiotic taxa.

Similarly,we performed this analysis on the entire dataset without removing the rare taxa. For this, our method selected 721 OTUs as abiotic taxa. The Bonferroni-corrected p-Value for this set is $6.94 \times 10^{-5}$ instead of 0.05. The p-values are in Dataset 6. Out of 721 OTUs, only 2 OTUs have p-value lower than the Bonferroni corrected p-value - OTU2709 (Order - Sphingobacteriales) and OTU2528 (Genus - Novosphingobium).

#### 2.7 Predicting enrichment rates from carbon catabolic phenotypes

To test if the carbon consumption capability can predict the enrichment rates of the bacterial isolates (Figure S17A), we regressed the enrichment rates on the carbon consumption capability. The carbon consumption ability was obtained from the growth measurements - whether or not an isolated strain could grow on a carbon source. As described in Section 1.10, maximum OD600 was used to obtain the maximum yield of the isolates on the different carbon sources. Strains that reached a maximum OD600 of 0.05 or beyond on a carbon source were designated as able to consume that carbon source; those that did not reach an OD600 of 0.05 on a given carbon source were designated as being unable to grow on that carbon source. The growth data were binarized in this way for the regression. This is the binary matrix in Figure S17A.

We have 21 isolates and 7 carbon sources. To prevent over-fitting, we used LASSO regularized linear regression to fit the model given by

$$r_{h,s} = \beta_0 + \sum_{j=1}^{7} \beta_j g_{j,s} + \epsilon_s \quad \text{(S5)}$$

where $g_{j,s} \in \{0,1\}$ represents the carbon utilization capability of isolate $s$ for carbon source $j$, $\beta_j$ are the regression coefficients for the carbon source $j$, $\beta_0$ is the intercept, and $\hat{r}_{h,s}$ represents the enrichment rate predicted by the LASSO regression, and $\epsilon_s$ is a noise term. The LASSO regression imposes a penalty on the regression coefficients (Figure S17B) $\beta_j$ by using the cost function to $\frac{1}{N}\|r_{h,s} - \hat{r}_{h,s}\|_2^2 + \lambda\|\beta\|_1$, where N is the number of samples, here the number of isolates and is equal to 21, and $\lambda$ is the regularization hyperparameter. The algorithm first optimizes $\lambda$ to find the lowest mean squared error in the predictions. This is done by iterated cross validation. The data are split into 4 folds. Holding one fold out, the regression is fit to the rest of the data for a range of $\lambda$ values. The coefficients obtained from the fit are used to predict the held out data, and the mean squared error is computed between the held out data and the prediction for each $\lambda$. This process is repeated by holding out each of the folds. This process is repeated 100 times, each time by splitting the data into different sets of 4 folds. This gives the mean squared error for the range of $\lambda$ values used. From this, the $\lambda$ corresponding to the minimum mean squared error is used as the optimal hyperparameter $\hat{\lambda}$. If the LASSO regression is able to find a sparse set of regression coefficients that predict the response variable well, there is a clear minimum in the mean squared error for a particular $\lambda$ as seen in the left hand column of Figure S16 [15]. If the regression fails, no clear minimum is obtained as seen in the right hand column of Figure S16.

We find that the regression is successful for the $\tau_{12}$ hybrid communities of Soil A and the $\tau_{12}$ and $\tau_9$ hybrid communities of Soil B, but fails for all others. For the control communities, naturally, $r_{c,s}$ is used as the enrichment rate instead of $r_{h,s}$. This indicates that when the algal impact on community assembly is strong, the carbon consumption capabilities of a bacterium can predict its enrichment rates, but in the control communities and the $\tau_3$ and $\tau_6$ hybrid communities where *C. reinhardtii* has little impact on community assembly, a strain's carbon consumption capabilities has no predictive power on the enrichment rates.

For each regression, LASSO estimates the regression coefficients $\beta_j$ which are the importance of each carbon source for predicting enrichment rates. For the cases where the regression is successful, we plot the regression coefficients corresponding to each carbon source in the top panels of Figure S17 C, D and E. For Soil B, Maltose, Ribose and Pyroglutamic acid have predictive power on the enrichment rates, and in Soil A, Maltose and Pyroglutamic acid have predictive power on the enrichment rates. It is noteworthy that glucose is not picked up as having predictive power in either of these cases. This result suggests that growth on carbon excreted by the algae is predictive of enrichment rates in the long growth period conditions.

To assess the out-of-sample predictive power of the model, we divided the data into test sets and training sets in numerous different combinations (4 fold split with 100 repeats). At the optimized penalty coefficient (hyperparameter), $\hat{\lambda}$, we fit the regression on the training set, obtain the regression coefficients and predict the test set. For each of these predictions, we obtained an $R^2$ score which determines the goodness of fit. The distribution of $R^2$ determined by cross-validation are plotted in the first column of the bottom panels of Figure S17C, D and E. To assess the significance of these values, we performed the regression again on synthetic data where the response variable (enrichment rates) were shuffled (randomized). For the shuffled data, we found the best penalty parameter $\hat{\lambda}$, and repeated the process just described to get the $R^2$ scores. We repeated this for 10 different shuffles, each with 4 fold splits and 10 repeats, to get a distribution of $R^2$ scores. This is plotted in the second column of the bottom panels of Figure S17C, D and E.

To compare the two distributions, we performed bootstrapping. We sampled the two distributions with replacement numerous times. Each time, we found the medians of the resulting distributions and subtracted them. This gave us a distribution of these differences. We tested for the null hypothesis that the median of the shuffled data is the same as or greater than the median of the actual data. In all cases, we found p-value $< 0.0001$, refuting the null hypothesis. This means that statistically, the regression on the actual data had significant predictive power compared to the null.

### 3 Notes on the datasets

#### 3.1 Dataset 1: Abundance data at ASV level

Here the 16S sequence counts for all the samples are presented. The rows represent the samples and columns represent the ASVs. The samples are named as follows: Soil Sample (A or B) Replicate (r1, r2, etc.) Community type (h for control communities, and hcr for hybrid communities) Growth Period (3d, 6d, 9d or 12d) Serial Dilution Round (r2, r4, r6, r8 or r10).

So, a sample with the name “Ar3hcr6dr6” represents soil sample A, replicate 3, hybrid community, 6 day growth period and dilution round 6. The name ”CR only” followed by a number indicates a sample with only *C. reinhardtii.* The numbers following the name represent the replicates. Cultures of only *C. reinhardtii* were sequenced to identify their contribution to the 16S sequence data.

#### 3.2 Dataset 2: Phylogeny of ASV data

Here the phylogeny of the 16S sequences are presented. The first column has the exact sequences, the next six columns present the phylogenetic information, and the final column maps the sequences and the phylogeny to the ASV number used in Dataset 1.

#### 3.3 Dataset 3: Abundance data at OTU level

Here the counts after grouping by OTUs is presented for all the samples are presented. The rows represent the samples and columns represent the OTUs. The naming convention of the samples is the same as in Dataset 1.

#### 3.4 Dataset 4: Phylogeny of OTU data

This table represents the taxonomy of the OTUs, including the feature ID and the confidence as given by the algorithm. The final column maps the phylogeny to the OTU number used in Dataset 1.

#### 3.5 Dataset 5: p-values from Fisher exact test of abiotic taxa

The p-values of the taxa contributing to displacement of the communities along PC1 for the analysis with the rare taxa removed, is presented here, according to the analysis described in Section 2.6 for both soil samples. The Bonferroni corrected p-value is also mentioned.

##### 3.6 Dataset 6: p-values from Fisher exact test of abiotic taxa, without removing the rare taxa

The p-values of the taxa contributing to displacement of the communities along PC1 for the analysis without removing the rare taxa, is presented here, according to the analysis described in Section 2.6 for both soil samples. The Bonferroni corrected p-value is also mentioned.

##### 3.7 Dataset 7: GC-MS data of algal spent media

The concentrations of the compounds secreted by *C. reinhardtii* on its own as measured by GC-MS are presented here. The concentrations of these compounds in the media is subtracted from measurement to assay for increasing or decreasing concentrations in time.

##### 3.8 Dataset 8: Sequence, phylogeny, abbreviation and origin of the bacterial isolates

For the 21 isolated bacterial strains, the sequence, taxonomy, abbreviation of the names used in Figure 6, and the community from which it was isolated are presented here. Note that for most of these isolates, we obtained multiple clones of the same isolate from different communities. The particular community from which the isolate was used for the experiments are presented here. The abbreviations used in Figure 6 and and Figure S17A are included.

##### 3.9 Dataset 9: Phylogenetic tree

The OTU data set with the rare taxa removed, as described in Methods, is used to create a newick tree file, which was used for Unifrac analysis.

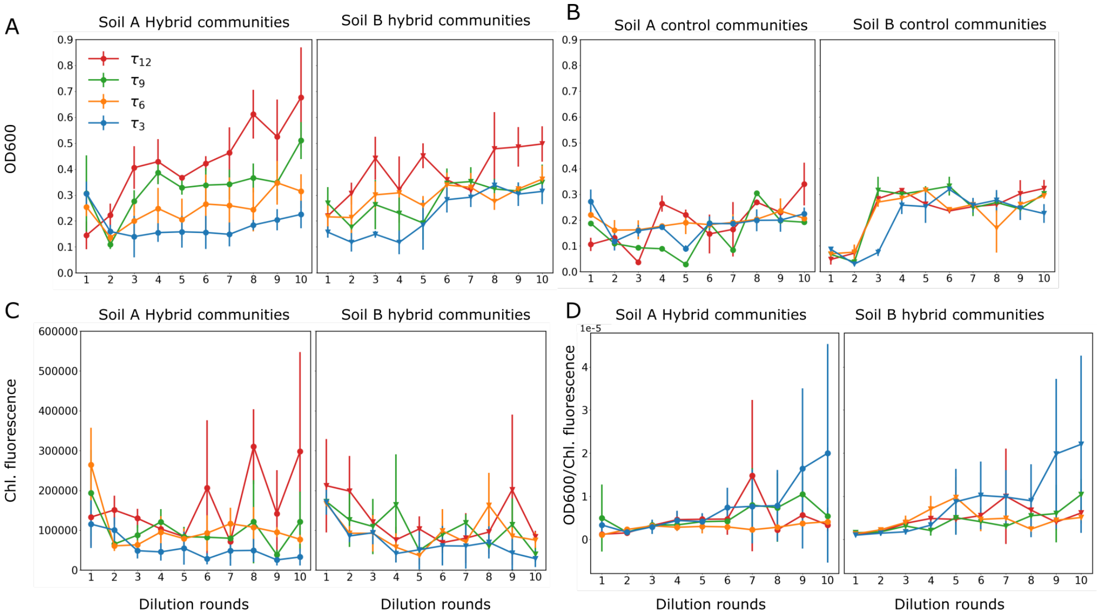

**Figure S1: OD600 and Chlorophyll fluorescence measurements:** The two columns in each panel correspond to the two soil samples. The error bars represent standard deviation over replicate communities. A shows the changes in OD600 after the end each dilution round for hybrid communities. B shows the changes in OD600 after the end each dilution round for control communities. C shows the changes in the auto-fluorescence of the chlorophyll content in the algae across dilution rounds for the hybrid communities. D shows the ratio of the above two quantities, i.e., OD 600:Chlorophyll content over dilution rounds for the hybrid communities.

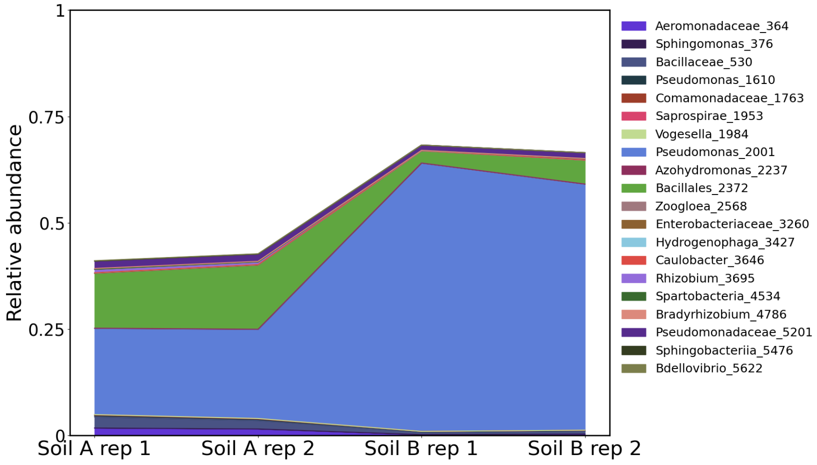

**Figure S2: Composition of initial soil inoculates:** The taxonomic composition of the initial soil inoculates, (after the treatment in the dark with drugs), with two replicates each, based on 16S sequencing is shown here. Only those OTUs that have a relative abundance greater than 20% at any time in any community are plotted. The relative abundance is plotted on the y-axis. The color map is the same as the color map of Figure 2 in the main text. The labels are the genus name of the OTU followed by the OTU number assigned during analysis. If the genus of a OTU is unknown, the next higher known classifier is used.

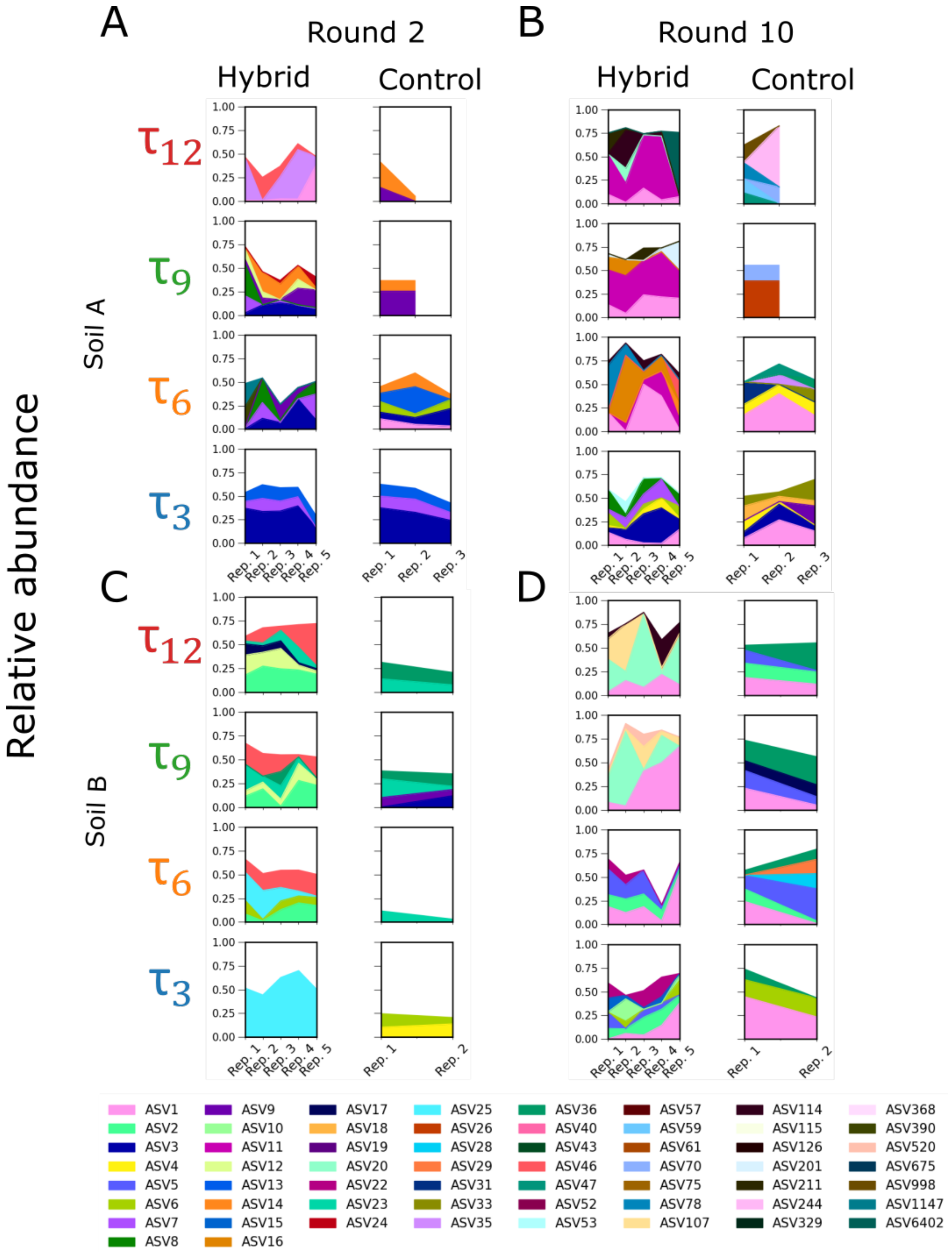

**Figure S3: Composition of the communities at the ASV level:** Same layout as Figure 2 of the main text. The taxonomic composition of the dominant taxa of the hybrid and control communities at the ASV level for serial dilution rounds 2 and 10 are shown here. By dominant taxa we mean those taxa that occur at a relative abundance of at least 20% in any community at any time point.

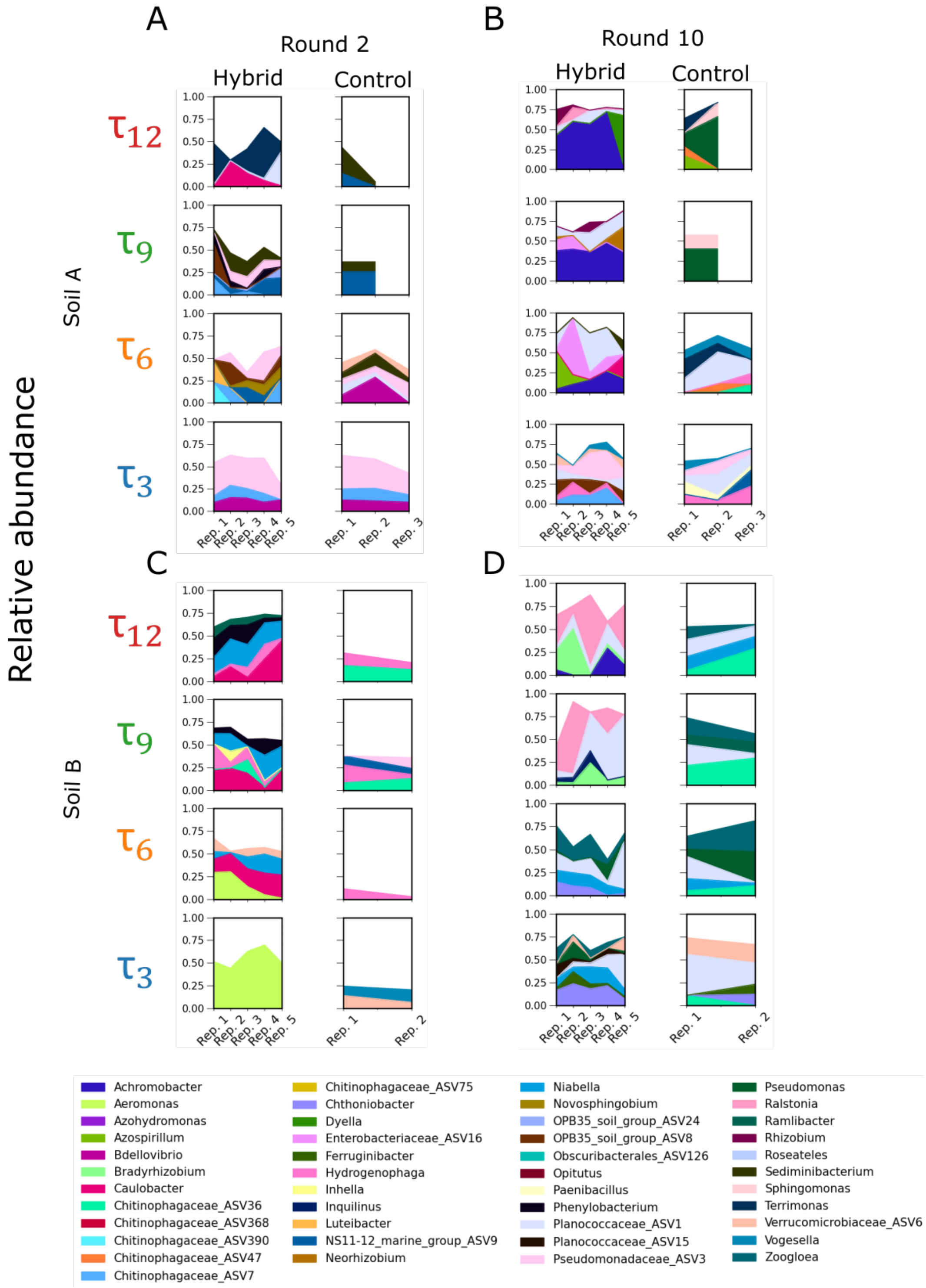

**Figure S4: Composition of the communities at the genus level:** Same layout as Figure 2 of the main text. The taxonomic composition of the dominant taxa of the hybrid and control communities at the genus level for serial dilution rounds 2 and 10 are shown here. By dominant taxa we mean those taxa that occur at a relative abundance of at least 20% in any community at any time point. Taxa with the same genus are grouped together. If the genus is not assigned, the next taxonomic classificiation is used along with the assigned ASV number.

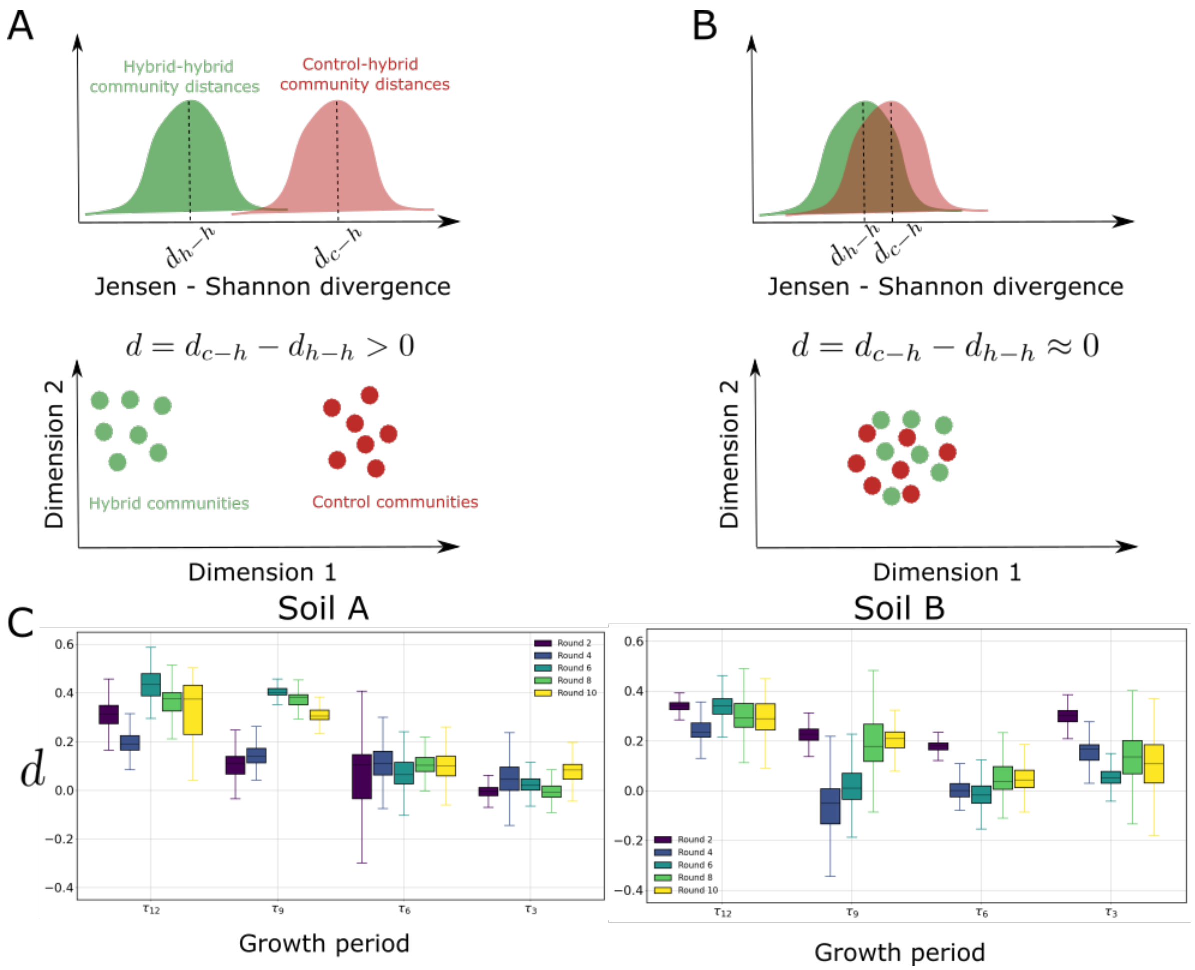

**Figure S5: Jensen-Shannon divergence between hybrid and control communities:** The Jensen-Shannon divergence (JSD) is computed between all pairs of communities. Then, the for each soil sample, for each dilution round, and for each growth period, the distribution of JSD is plotted for hybrid - hybrid communities and control-hybrid communities. A and B show hypothetic sketches of what these two distributions of JSD might look like. A shows the case where hybrid communities are taxonomically more similar to each other than they are to the control communities because the median distance between hybrid communities ($d_{h-h}$) is smaller than the median distance between control and hybrid communities ($d_{c-h}$), and $d = d_{c-h} - d_{h-h}$ is positive. Such a situation could be represented in the bottom row of A. If the distributions are as sketched in B, then the hybrid communities are taxonomically similar to the control communities because the median distance between hybrid communities ($d_{h-h}$) is similar to the median distance between control and hybrid communities ($d_{c-h}$), and $d = d_{c-h} - d_{h-h}$ is roughly zero. Such a situation would look like the sketch in the bottom row of B. In C, $d$ is computed across all replicates and plotted on the y-axis for each dilution round and growth period plotted along the x-axis. By dilution round 10, in both soil samples $d$ is significantly positive for the $\tau_{12}$ and $\tau_9$ communities (p-values $< 0.002$, testing for the null hypothesis that $d$ is equal to or less than zero.) However, the $\tau_3$ communities do not have a significantly positive $d$ (p-values 0.13 and 0.07).

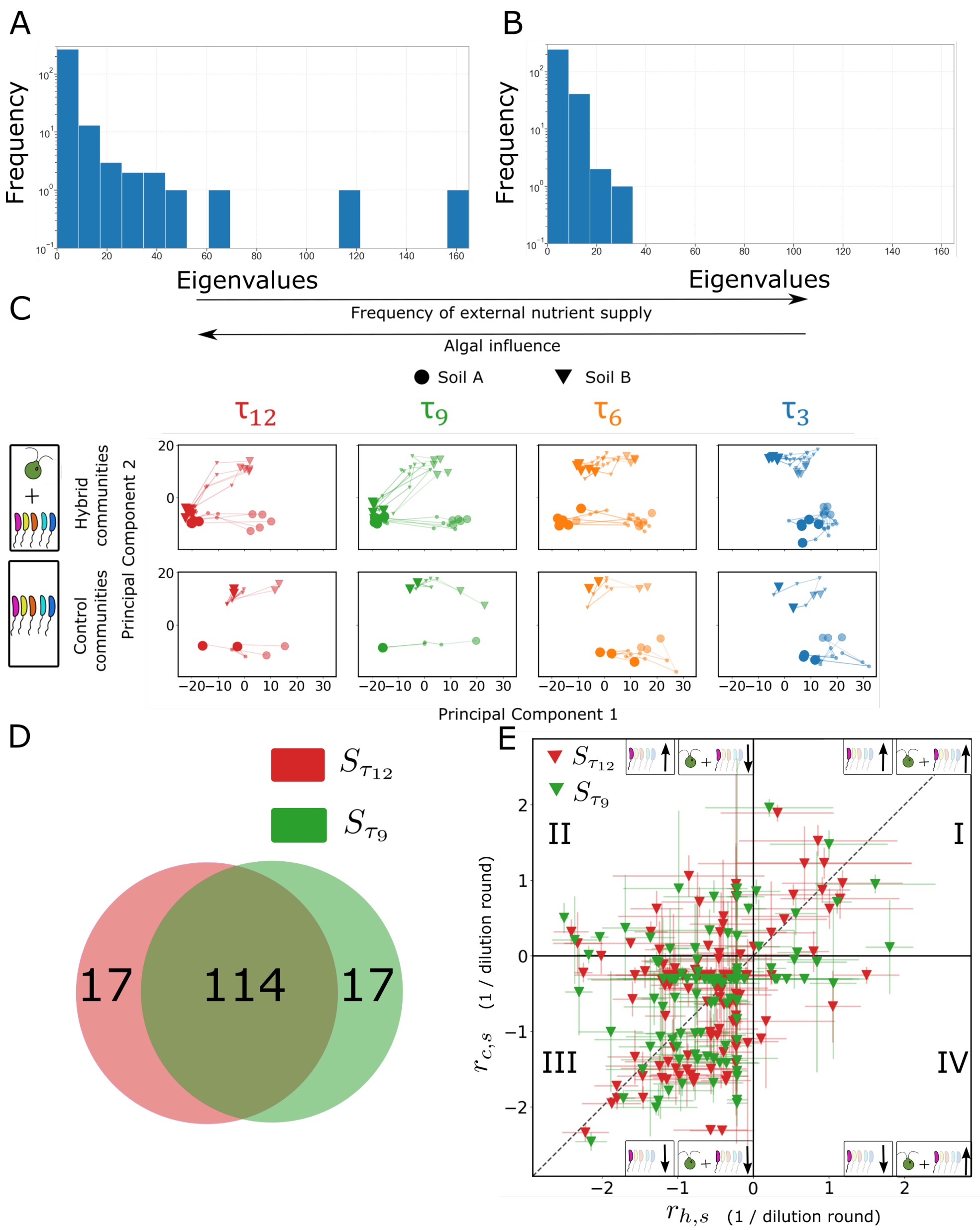

**Figure S6: Principle components analysis without removal of rare taxa:** PCA was performed on the entire data set without any removal of rare taxa. A shows the distribution of eigenvalues for the data, showing two modes with high eigenvalues. B shows the distribution of eigenvalues for shuffled data. The two dominant modes are absent. C shows the PCA results on the entire data without removal of rare taxa. The circles and triangles differentiate the two soil samples. The different colors represent the different growth periods. The intermediate translucent markers represent the initial round of serial dilution, the large solid markers represent the final round of serial dilution. The smallest markers represent the intermediate rounds of serial dilution. The results are very similar to Figure 4 in the main text, showing that the two soil samples converge for hybrid $\tau_9$ and and $\tau_{12}$ communities, but not for the control or the $\tau_3$ and $\tau_6$ communities. D shows the intersection of the biotic taxa, i.e. those that contribute to the motion along PC2 for $\tau_{12}$ ($S_{\tau_{12}}$) and $\tau_9$ ($S_{\tau_9}$) hybrid communities of soil sample B. As with thresholded data (Figure 5C), majority of the taxa that cause displacement along PC2 are common to the two growth periods. E shows the enrichment plot of these biotic taxa. As for the analysis with rare taxa removed, (Figure 5E, main text), most of these taxa are on the left half of the figure.

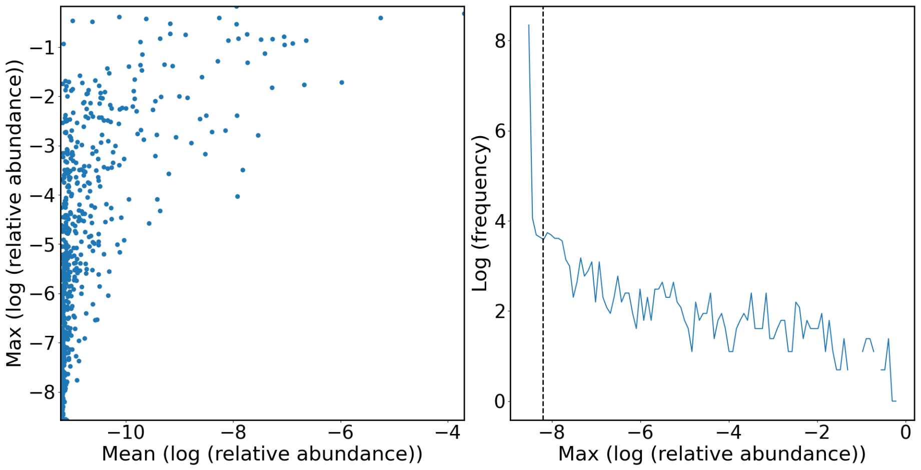

**Figure S7: Selecting the threshold for removal of OTUs for PCA analysis:** We computed the maximum and mean of logarithm of the relative abundance of all taxa across all dilution rounds and communities. In the left panel this maximum and mean are plotted for all the taxa. We note that majority of the taxa have a very low mean and maximum relative abundance across all samples and all times, indicating they are rare in all communities at all times. To set a cut-off, we plot the frequency of the maximum of the logarithm of the relative abundance across all dilution rounds and for all samples for each OTU in the right panel. The sharp rise in frequency below -8 shows that, as expected, many taxa have a low maximum relative abundance across all time points and samples. Based on this curve, the cutoff is set at a maximum logarithm of relative abundance of -8.2 indicated by the vertical dashed line in the right panel. Out of 5236 OTUs, this process removed 4360 OTUs.

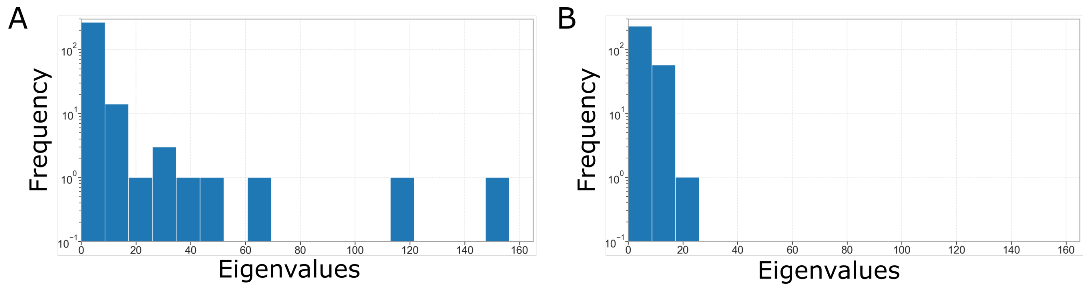

**Figure S8: Eigenvalue distributions after removing rare taxa:** PCA is performed on the data with the rare OTUs remove, and the frequency of the eigenvalues are plotted in A. The distribution shows that there are two modes with high eigenvalues that are distinct from the rest. Panel B shows the frequency distribution of the eigenvalues for the shuffled data. Here the two modes with the highest eigenvalues are absent.

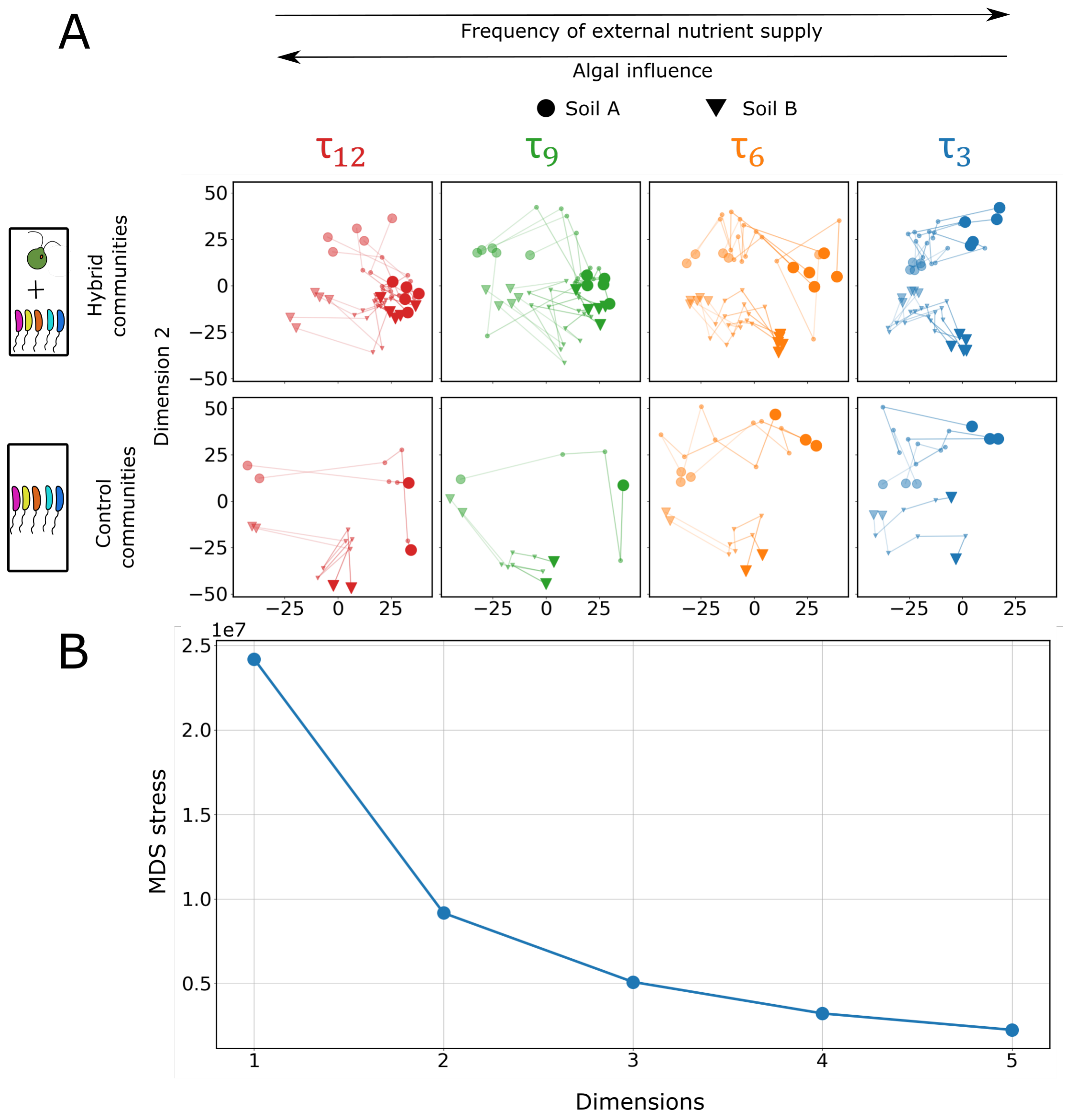

**Figure S9: MDS embedding of Aitchison's distances:** Aitchison's distances were computed using the entire data set between all pairs of communities. These distances were then embedded using the metric Multi Dimensional Scaling (MDS, using the MDS function scikit-learn's manifold class) using Euclidean distances in two dimensions, shown in A. The circles and triangles differentiate the two soil samples. The different colors represent the different growth periods. The intermediate translucent markers represent the initial round of serial dilution, the large solid markers represent the final round of serial dilution. The smallest markers represent the intermediate rounds of serial dilution. As with the PCA analysis (Figure 4), we observe that the $\tau_{12}$ and $\tau_9$ hybrid communities converge by dilution round 10, whereas the $\tau_3$ and $\tau_6$ hybrid communities and none of the control communities converge. B shows the stress of the MDS embedding.

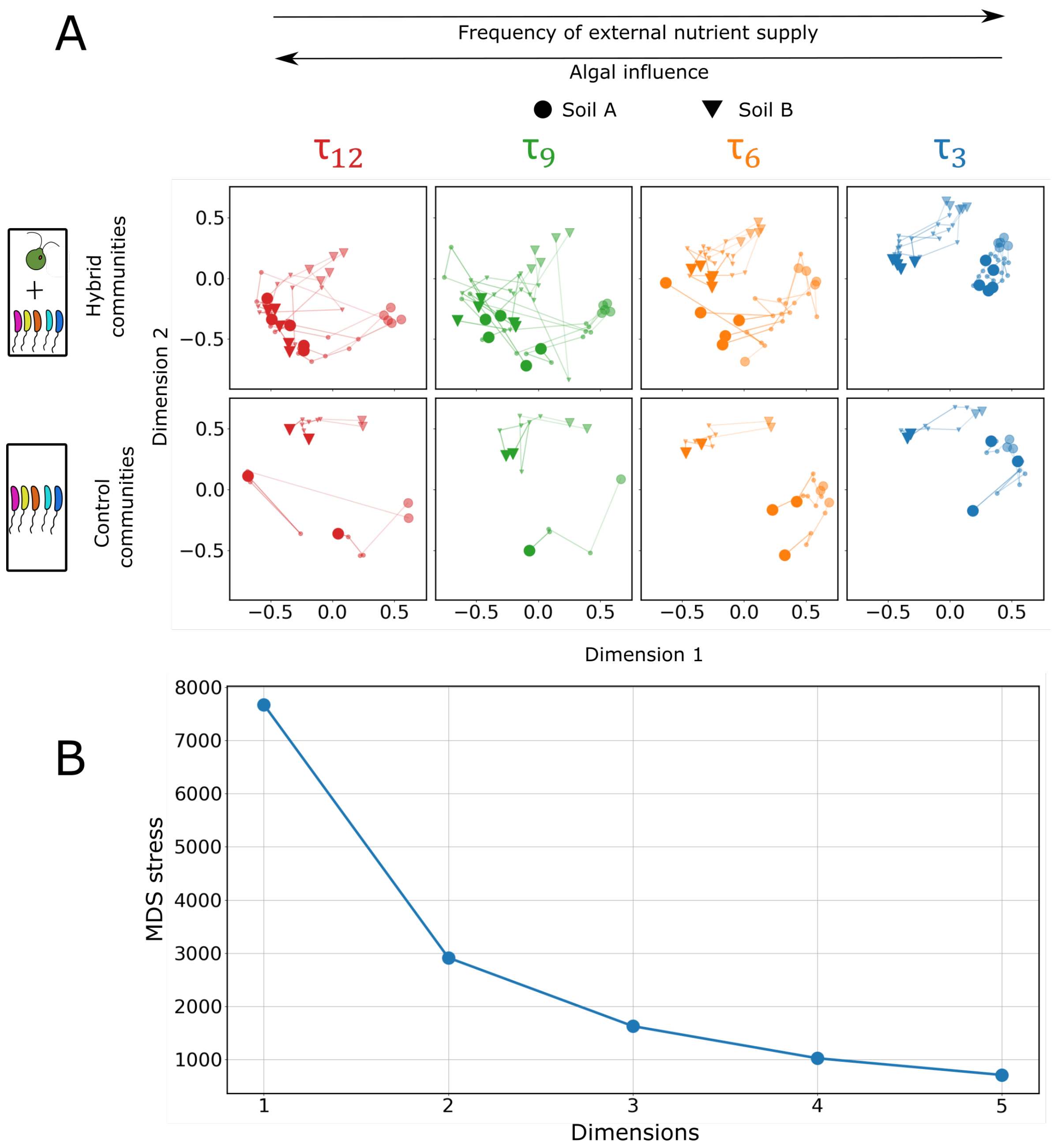

**Figure S10: MDS embedding of the Unifrac distances:** Unifrac distances were computed on the data set with rare OTUs removed using a phylogenetic tree, between all pairs of communities. These distances were then embedded using the metric Multi Dimensional Scaling (MDS, using the MDS function scikit-learn's manifold class) using Euclidean distances in two dimensions, shown in A. The circles and triangles differentiate the two soil samples. The different colors represent the different growth periods. The intermediate translucent markers represent the initial round of serial dilution, the large solid markers represent the final round of serial dilution. The smallest markers represent the intermediate rounds of serial dilution. As with the PCA analysis (Figure 4, main text), we observe that the $\tau_{12}$ and $\tau_9$ hybrid communities converge by dilution round 10, whereas the $\tau_3$ and $\tau_6$ hybrid communities and none of the control communities converge. B shows the stress of the MDS embedding.

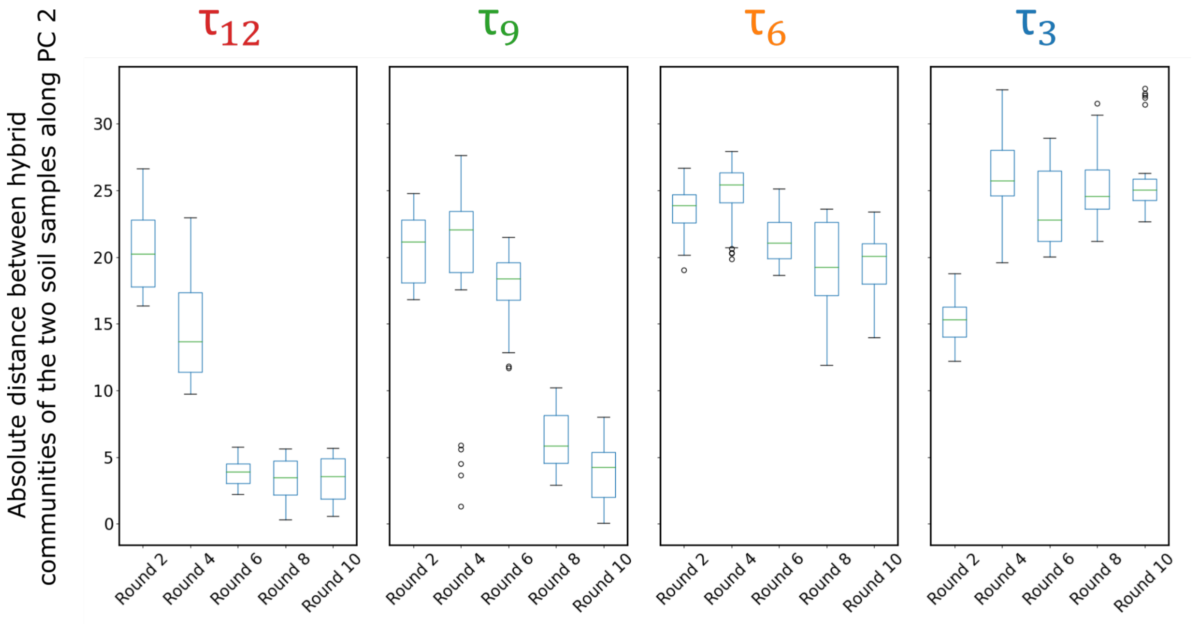

**Figure S11: Distance between $\tau_3$ hybrid communities of the two soil samples does not decrease:** The absolute distance between the hybrid communities along PC2 are plotted for the four growth periods. For $\tau_{12}$ and $\tau_9$ communities, the distance decreases, signifying convergence, while for $\tau_3$ communities, the distance increases over dilution rounds.

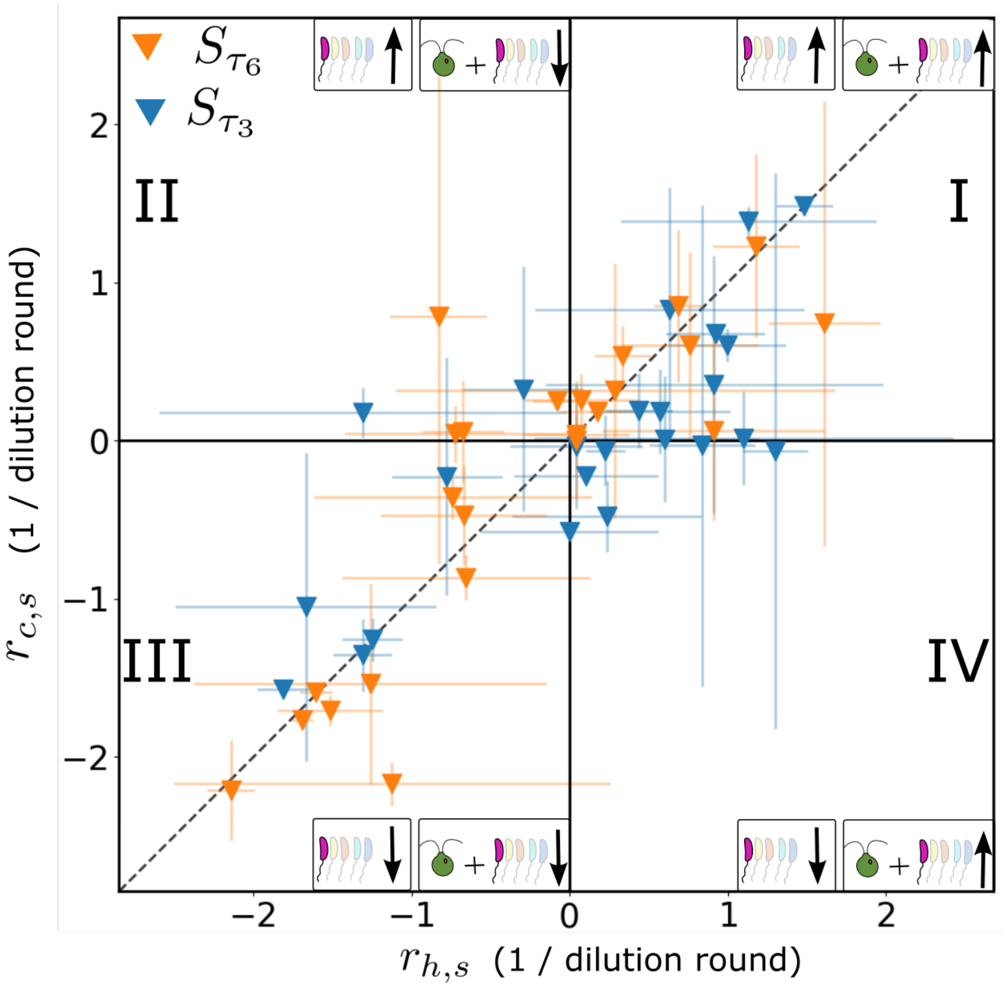

**Figure S12: Enrichment plot for biotic taxa in $\tau_6$ and $\tau_3$ hybrid communities:** The construction of this plot is identical to Figure 5D of the main text. For the biotic taxa selected for their contribution to movement along PC2 in the $\tau_{12}$ and $\tau_9$ hybrid communities of soil sample B, their enrichment in the $\tau_6$ and $\tau_3$ hybrid communities are computed as described in the main text and plotted here. As opposed to the $\tau_9$ and $\tau_{12}$ hybrid communities (Figure 5D), in the $\tau_3$ and $\tau_6$ hybrid communities, these taxa dominantly lie on the diagonal, where the presence of *C. reinhardtii* has no significant effect on the enrichment dynamics.

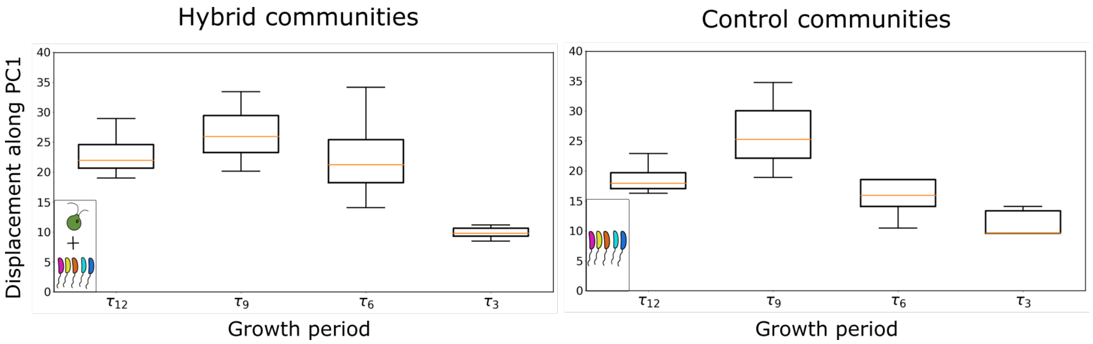

**Figure S13: Displacement along PC1 depends on the growth period:** Exogenous nutrient supply rate, which is equivalent to the growth period and is the experimentally controlled abiotic factor, controls the displacement along PC1, the abiotic axis. Communities for which the exogenous nutrients are supplied very frequently, move shorter distances along PC1 compared to the communities where the exogenous nutrient supply is infrequent. This is true for both hybrid (left panel) and control (right panel) communities, for both soil samples. With the null hypothesis that the distribution of displacements laong PC1 is the same for the $\tau_{12}$ and $\tau_3$ communities, the Kolmogorov Smirnoff test gives a p-value of $2 \times 10^{-8}$, indicating that the null hypothesis is false.

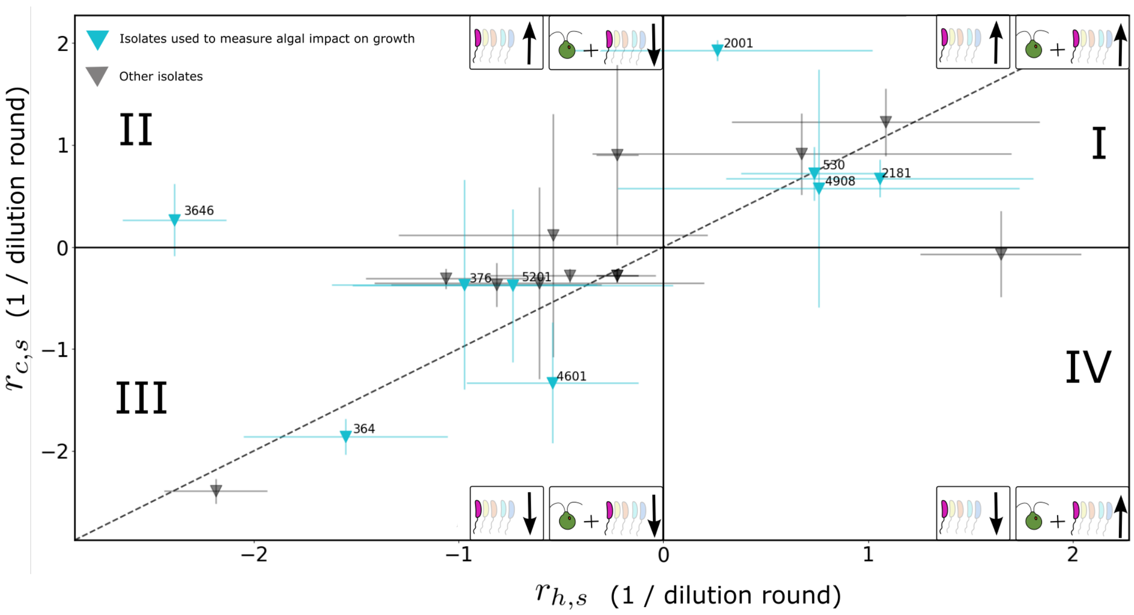

**Figure S14: Enrichment plot for all isolates:** For the 21 isolated bacterial strains, their enrichment in the control and hybrid communities, averaged over the $\tau_9$ and $\tau_{12}$ growth periods are plotted. The error bars indicate standard deviation. The isolates span all four quadrants of the plot. The isolates in blue are used in the measurement of algal impact on growth in Figure 6D. For visual clarity, only their assigned OTU numbers are shown.

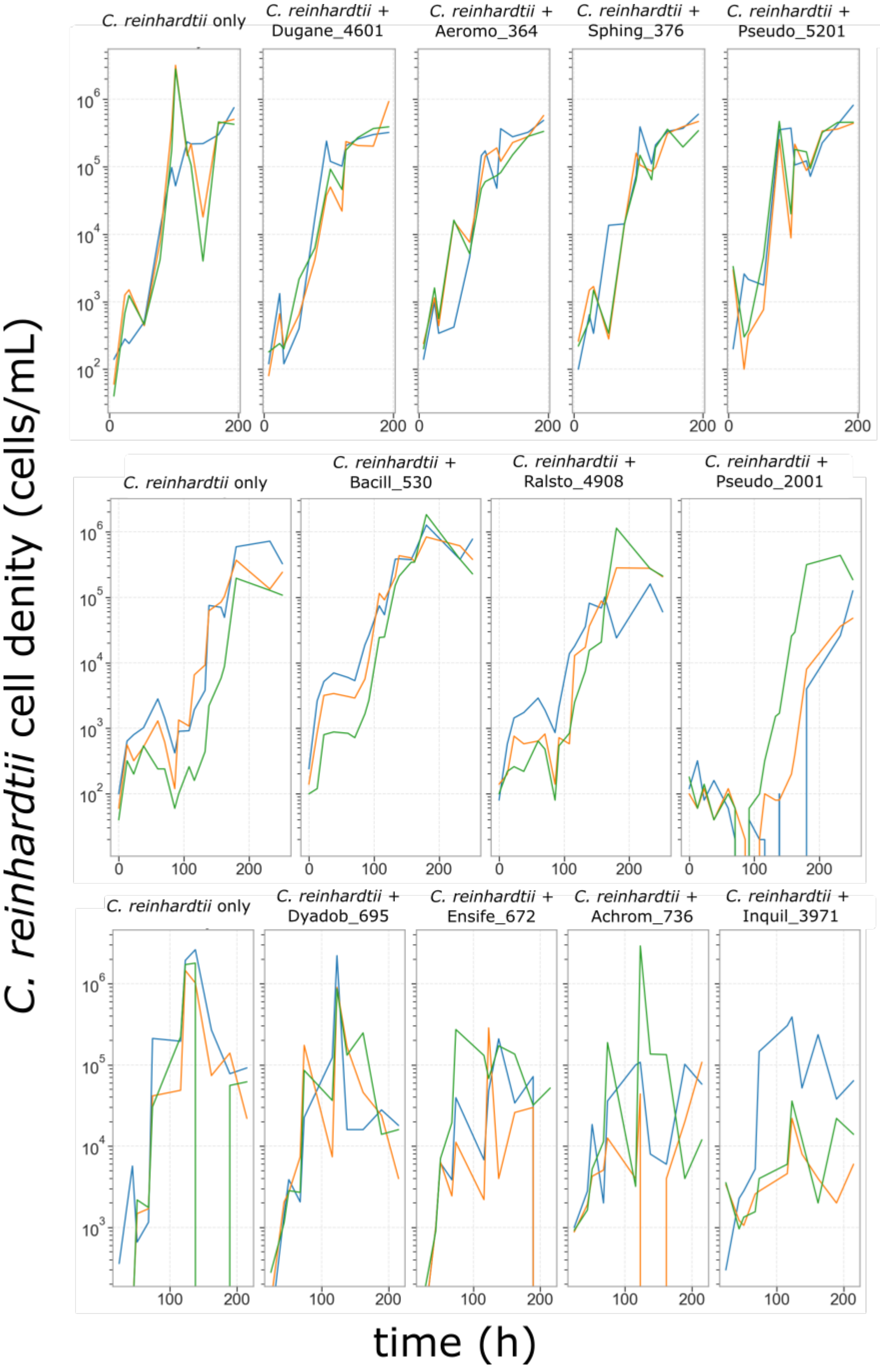

**Figure S15: Raw data from flow cytometry:** The raw data from flow cytometry is plotted here. There were 3 runs in total, represented by each row in the figure. For each run, we had the control of *C. reinhardtii* growing on their own. There were three replicates for each measurement, represented by the three curves in each panel.

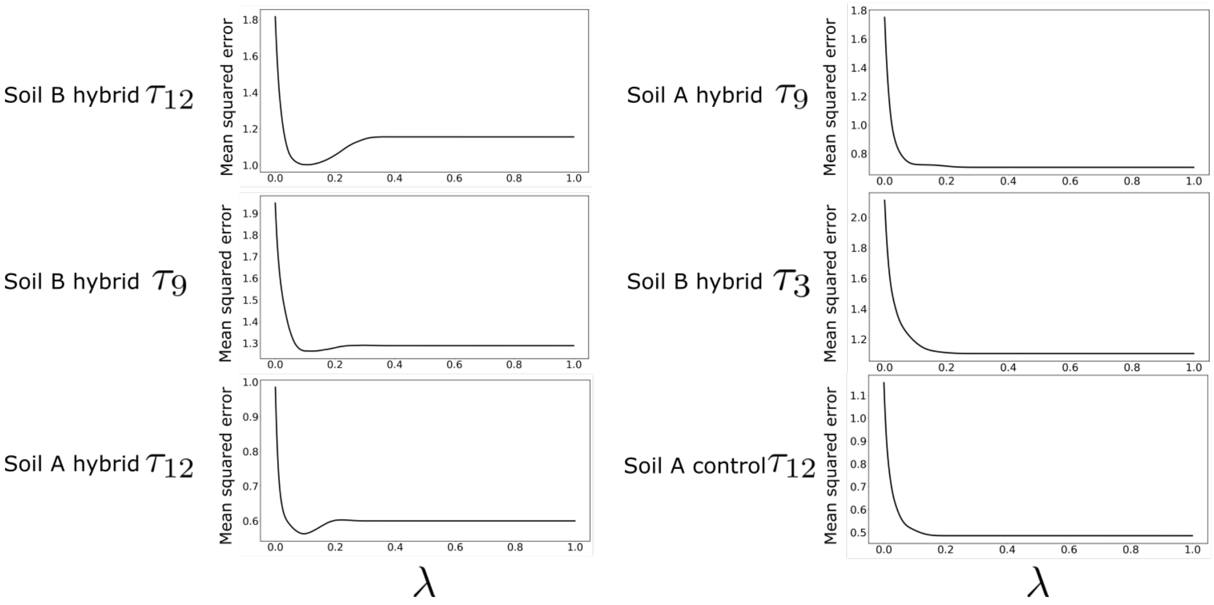

**Figure S16: Determination of optimal hyperparameter ($\lambda$) for LASSO regression via cross-validation:** Mean squared error in held out data determined by iterated 4-fold cross validation for a range of $\lambda$. Here, in the first column, we show the cases where the LASSO regression finds an optimal penalty coefficient $\lambda$: there is a well defined minimum in the mean squared error for a particular $\lambda$. In the second column, we show that the LASSO regression fails to find a minimum for $\lambda$, because there is no well defined minimum for the mean squared error for a range of $\lambda$. Therefore the regression fails in these cases, and as shown in[15], in such cases, the LASSO regression and only returns an intercept and there is likely no sparse solution to the regression problem.

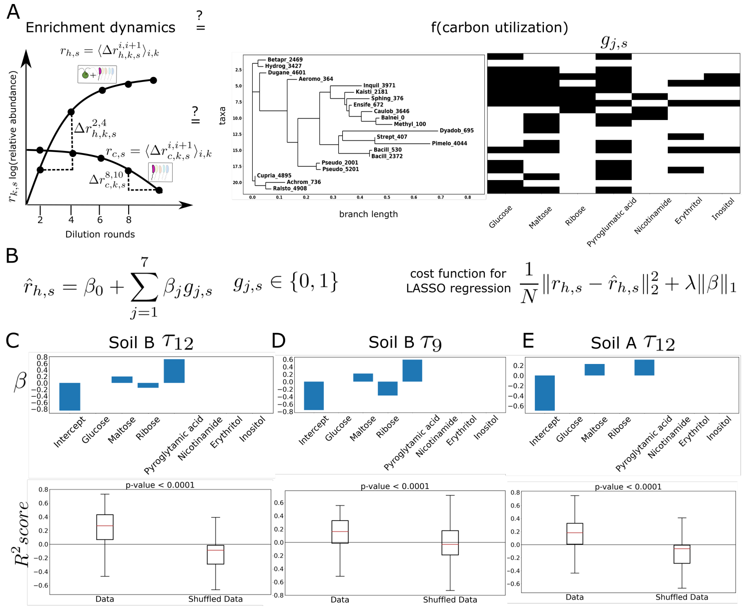

**Figure S17: Carbon catabolism weakly predicts enrichment rates in long growth period hybrid communities:** A. We ask if the enrichment rates ($r_{h,s}$, $r_{c,s}$) of the bacterial isolated strains are related to their ability to grow on a carbon source, represented here as a binary matrix $g_{j,s}$ for the isolated strains $s$ and carbon source $j$. The isolated strains are ordered according to their position on a phylogenetic tree. The naming convention is first six letters of their genus followed by the assigned number. If the genus is not assigned, the next assigned taxonomy is used. The binary matrix to the right indicates growth or no growth on each carbon source as measured in a plate reader experiment (see Sections 1.10 and 2.7). B. The enrichment rate of the isolate $s$ is predicted ($\hat{r}_{h,s}$) through a regression against its carbon consumption ability $g_{j,s} \in 0, 1$, with coefficients $\beta_j$ and intercept $\beta_0$. Regularized, LASSO regression is performed, with the cost function as shown, where N is the number of samples (here, 21 isolates), $r_{h,s}$ the true enrichment, $\hat{r}_{h,s}$ the predicted enrichment, $\lambda$ is the penalty coefficient and $\beta$ is the vector of the regression coefficients $\beta_j$ and $\beta_0$. The optimal value of $\lambda$ is found first by iterated cross-validation with a 4 fold split and 100 repeats, and then the regression coefficients are determined at the optimal hyperparameter $\hat{\lambda}$ (see Section 2.7 and Figure S16) C. The top panel shows the regression coefficients $\beta_j$ and the intercept $\beta_0$ for enrichment rates for isolates in Soil B $\tau_{12}$ hybrid communities. The LASSO regression picks Maltose, Ribose and Pyroglutamic acid as the carbon sources with non-zero regression coefficients. To test if the regression has out of sample predictive power, we train the regression on part of the data at the optimal hyperparameter $\hat{\lambda}$, and test its prediction on the rest. We report an $R^2$ on the held out data to measure the out-of-sample predictive power. The bottom panel shows the distribution of this $R^2$ score on the data and on the data obtained by shuffling the enrichment rates. A bootstrap test is performed with the null hypothesis that the median of the $R^2$ score of the shuffled data is larger than or equal to that of the actual data. The low p-value suggests that this null is false, and that the median of the $R^2$ distribution of the actual data is higher than that of the shuffled data, indicating that the LASSO regression is significantly predictive on the data. D. The top panel shows the regression coefficients $\beta_j$ and the intercept $\beta_0$ for Soil B $\tau_9$ hybrid communities. The regression picks Maltose, Ribose and Pyroglutamic acid as the carbon sources with the predictive power. The distribution at the bottom panel of the $R^2$ scores for the data and the shuffled data shows that LASSO regression has a significant predictive power on the data. E. The top panel shows the regression coefficients $\beta_j$ and the intercept $\beta_0$ for Soil A $\tau_{12}$ hybrid communities. The LASSO regression picks Maltose and Pyroglutamic acid as the carbon sources with the predictive power. The distribution at the bottom panel of the $R^2$ scores for the data and the shuffled data shows that the regression has significant out-of-sample predictive power.

| Compound | Concentration |
|---|---|
| $C_6H_{12}O_6$ (glucose) | 1.333 mM |
| $NH_4Cl$ | 8 mM |
| $KH_2PO_4$, $K_2HPO_4$ | 3.1 mM |
| $MgSO_4$ | 0.1 mM |
| $CaCl_2$ | 1 mM |
| $C_{10}H_{16}N_2O_8$ (EDTA) | 5.5 $\mu$M |
| $FeSO_4$ | 5.5 $\mu$M |
| $H_3BO_4$ | 15 $\mu$M |
| $ZnSO_4$ | 0.5 $\mu$M |
| $MnCl_2$ | 3.5 $\mu$M |
| $Na_2MoO_4$ | 0.58 $\mu$M |
| $CuSO_4$ | 0.15 $\mu$M |
| $Co(NO_3)_2$ | 0.8 $\mu$M |
| NaOH | 999 $\mu$M |
| $FeSO_4 \cdot 7H_2O$ | 999 $\mu$M |
| NaCl | 999 $\mu$M |

**Table S1:** Modified 1/2x Taub medium composition

| Compound | Concentration |
|---|---|
| Tris-Cl | 20 mM |
| Sodium EDTA | 2 mM |
| Triton X-100 | 1.2% |
| Lysozyme from chicken egg | 20 mg/mL |

**Table S2:** Lysis buffer composition for DNA extraction

| Reagent | Volume |
|---|---|
| PCR grade water | 13 $\mu$L |
| Forward primer (10 $\mu$M) | 0.5 $\mu$L |
| Reverse primer (10 $\mu$M) | 0.5 $\mu$L |
| Template DNA | 1 $\mu$L |
| Platinum Hot Start Master Mix (2X) | 10 $\mu$L |

**Table S3:** Reagents for PCR for EMP protocol

| Temperature | Time | Repeat |
|---|---|---|
| 94 C | 3 min | |
| 94 C | 45 s | 45x |
| 50 C | 60 s | 45x |
| 72 C | 90 s | 45x |
| 72 C | 10 min | |
| 4 C | hold | |

**Table S4:** Thermocycler settings for EMP protocol

| Reagent | Volume |
|---|---|
| PCR grade water | 7 $\mu$L |
| Forward primer (10 $\mu$M) | 1 $\mu$L |
| Reverse primer (10 $\mu$M) | 1 $\mu$L |
| Cell suspension | 1 $\mu$L |
| Platinum Hot Start Master Mix (2X) | 10 $\mu$L |

**Table S5:** Reagents for colony PCR

| Temperature | Time | Repeat |
|---|---|---|
| 98 C | 3 min | |
| 98 C | 10 s | 40x |
| 55 C | 30 s | 40x |
| 72 C | 45 s | 40x |
| 72 C | 10 min | |
| 4 C | hold | |

**Table S6:** Thermocycler settings for colony PCR

| Compound excreted by *C. reinhardtii* in significant quantities |
|---|
| 2-O-Glycerol-$\alpha$-d-galactopyranoside |
| Digalactosylglycerol |
| Erythritol |
| Galacotse |
| Glyceric acid |
| Inositol,myo |
| Malic acid |
| Maltose |
| Nicotinamide |
| Proline |
| Putrescine |
| Pyroglutamic acid |
| Ribose |
| Threitol |
| Threonic acid |

**Table S7:** Compounds excreted in significant amounts by *C. reinhardtii* grown on its own
